# First gene-edited calf with reduced susceptibility to a major viral pathogen

**DOI:** 10.1101/2022.12.08.519336

**Authors:** Aspen M Workman, Michael P Heaton, Brian L Vander Ley, Dennis A Webster, Luke Sherry, Sabreena Larson, Theodore S Kalbfleisch, Gregory P Harhay, Erin E Jobman, Daniel F Carlson, Tad S Sonstegard

**Affiliations:** US Meat Animal Research Center, United States Department of Agriculture (USDA), Agricultural Research Service (ARS), Clay Center, NE, 68933, USA; University of Nebraska-Lincoln, Great Plains Veterinary Educational Center, Clay Center, Nebraska, 68933, USA; Recombinetics Inc., Eagan, MN, 55121, USA; Acceligen Inc., Eagan, MN, 55121, USA; Department of Veterinary Science, Gluck Equine Research Center, University of Kentucky, Lexington, KY, 40506, USA

**Author notes:** Corresponding author: Aspen Workman, US Meat Animal Research Center, 844 Road 313, Spur 18 D, Clay Center, NE 68933.

## Abstract

Bovine viral diarrhea virus (BVDV) is one of the most important viruses affecting the health and well-being of bovine species throughout the world. Here we used CRISPR-mediated homology-directed repair and somatic cell nuclear transfer to produce a live calf with a six amino acid substitution in the BVDV binding domain of bovine CD46. The result was a gene-edited calf with dramatically reduced susceptibility to infection as measured by clinical signs and the lack of viral infection in white blood cells. The edited calf has no off-target edits and appears normal and healthy at 16 months of age without obvious adverse effects from the on-target edit. This precision bred, proof-of-concept animal provides the first evidence that intentional genome alterations in CD46 may reduce the burden of BVDV-associated diseases in cattle, and is consistent with our stepwise, *in vitro* and *ex vivo* experiments with cell lines and matched fetal clones.

## Introduction

Bovine viral diarrhea virus (BVDV, family *Flaviviridae*, genus *Pestivirus*) is a ubiquitous pathogen of cattle that is associated with gastrointestinal and respiratory diseases and reproductive failure in cattle worldwide^1,2^. Following exposure, BVDV replicates in the oronasal mucosa as well as the tonsil before spreading to the regional lymph nodes^3,4^. BVDV infection of lymphocyte and monocyte populations results in immune suppression and systemic dissemination of the virus to multiple organs, including the digestive and reproductive tracts^2^. In pregnant cattle, BVDV crosses the placenta and infects the fetus resulting in abortion, congenital malformations, or birth of immunotolerant and persistently infected (BVDV-PI) calves^5^. While vaccines for BVDV have been available for more than 50 years, the extensive genetic and antigenic diversity in circulating field strains of BVDV pose a challenge to making these vaccines broadly protective^6^. Thus, if available, cattle with reduced genetic susceptibility to BVDV would complement current strategies to control BVDV-associated diseases.

Modifying pathogen receptors via genome-editing offers an innovative way to control viral infections in livestock^7–9^. Bovine CD46 is the main cellular receptor for BVDV,^10^ and CRISPR/Cas9 mediated knockout of CD46 in Madin-Darby bovine kidney (MDBK) cells results in a significant reduction in BVDV susceptibility *in vitro*^11–13^. Yet, it is unknown whether editing CD46 will reduce an animal’s susceptibility to BVDV *in vivo*. Like many viruses grown in cell culture, BVDV has been reported to rapidly adapt *in vitro* to infect cells lacking CD46 by acquiring mutations that enhance heparan sulfate mediated binding and entry^12^. However, viral adaptations to host receptors occurring *in vitro* do not necessarily correlate with *in vivo* outcomes^14,15^. Thus, the success of a gene-editing strategy for reducing viral infection can be difficult to accurately predict.

Deleting the entire *CD46* gene *in vivo* would likely have severe deleterious consequences. In most mammals, CD46 is ubiquitously expressed on all nucleated cells and is critical in several diverse biological processes^16^. The essential roles for CD46 function include complement inactivation, modulation of T-cell activation, and fertility^17,18^. For these reasons, minimal and precise genomic alterations are necessary if the normal cellular functions of CD46 are to be preserved in the host animal. Important work by Krey *et al*. mapped the BVDV receptor-binding sites of CD46 to two peptide domains, E_66_QIV_69_ and G_82_QVLAL_87_, which are located on antiparallel beta sheets in the amino terminus of the protein^10^. Further, by expressing chimeric CD46 molecules in porcine cells, they showed that replacing the bovine CD46 G_82_QVLAL_87_ residues with ALPTFS prevented BVDV cellular entry and blocked infection *in vitro*^10^.

Building on their discovery, our goal was to use gene-editing via homology directed repair to make a precise 18-nt replacement in the endogenous *CD46* gene to produce cells expressing homozygous CD46 with the A_82_LPTFS_87_ substitution in the BVDV binding domain. In stepwise experiments we measured the impact of this edit on BVDV susceptibility in immortalized cells, primary cells from fetal tissues, cells from a live CD46-edited calf, and in a natural exposure challenge study with the same edited calf.

## Materials and Methods

### Ethics Statement

All protocols for reproductive cloning, fetal tissue collection, and birthing were reviewed and approved by the Institutional Animal Care and Use Committee (IACUC) of TransOva Genetics (IACUC Project ID 68-000176-00). All protocols for the gene-edited calf and the BVDV challenge study were reviewed and approved by the IACUC of the University of Nebraska–Lincoln, an AAALAC International Accredited institution (IACUC Project ID 2111).

### Protein folding prediction of CD46 extracellular domains

The predicted protein structure of the CD46 extracellular domain, minus the signal peptide, was compared with the wild-type G_82_QVLAL_87_ residues and the substituted A_82_LPTFS_87_ residues. These extracellular domains consisted of residues 43-313 of the National Center for Biotechnology Information (NCBI) reference sequence: NP_898903.2. An artificial general intelligence (AGI) software that uses machine learning was used to fold the CD46 extracellular domain variants (AlphaFold v2.2.2)^19,20^. Each of the two CD46 variants were folded separately and the .pdb structure file with the highest confidence was selected as the best representation of the biologically relevant structure. The two CD46 variant files were loaded into a molecular graphics system (PyMOL Molecular Graphics System v2.5.3; Schrödinger, LLC, New York, NY), aligned, and rotated to view the variant region and the BVDV binding platform.

### CRISPR/Cas9 editing of CD46 in MDBK cells and Gir fibroblasts

BVDV-free Madin-Darby bovine kidney cells (MDBK; ATCC CCL-22) were obtained from the American Type Culture Collection (ATCC; Rockville, MD). Primary skin fibroblasts used for reproductive cloning were from a Gir breed (Bos indicus) female (TO470).

#### Gene editing and detection reagents

CRISPR/Cas9 technology was used to create altered alleles of CD46 in both MDBK and primary fibroblast cells. For the CD46 gene deletion (CD46Δ), the entire open reading frame of CD46 was removed, while for CD46 A_82_LPTFS_87_, six amino acids were substituted. The combination of cell type, guide RNA, and homology dependent repair templates for each edit are listed in **Table 1** according to cell type and gene editing outcome desired. Individual gRNAs were designed using Cas-Designer ^21^ and selected based on proximity to the target sequence and fewest potential off-target sites.

**Table 1.**
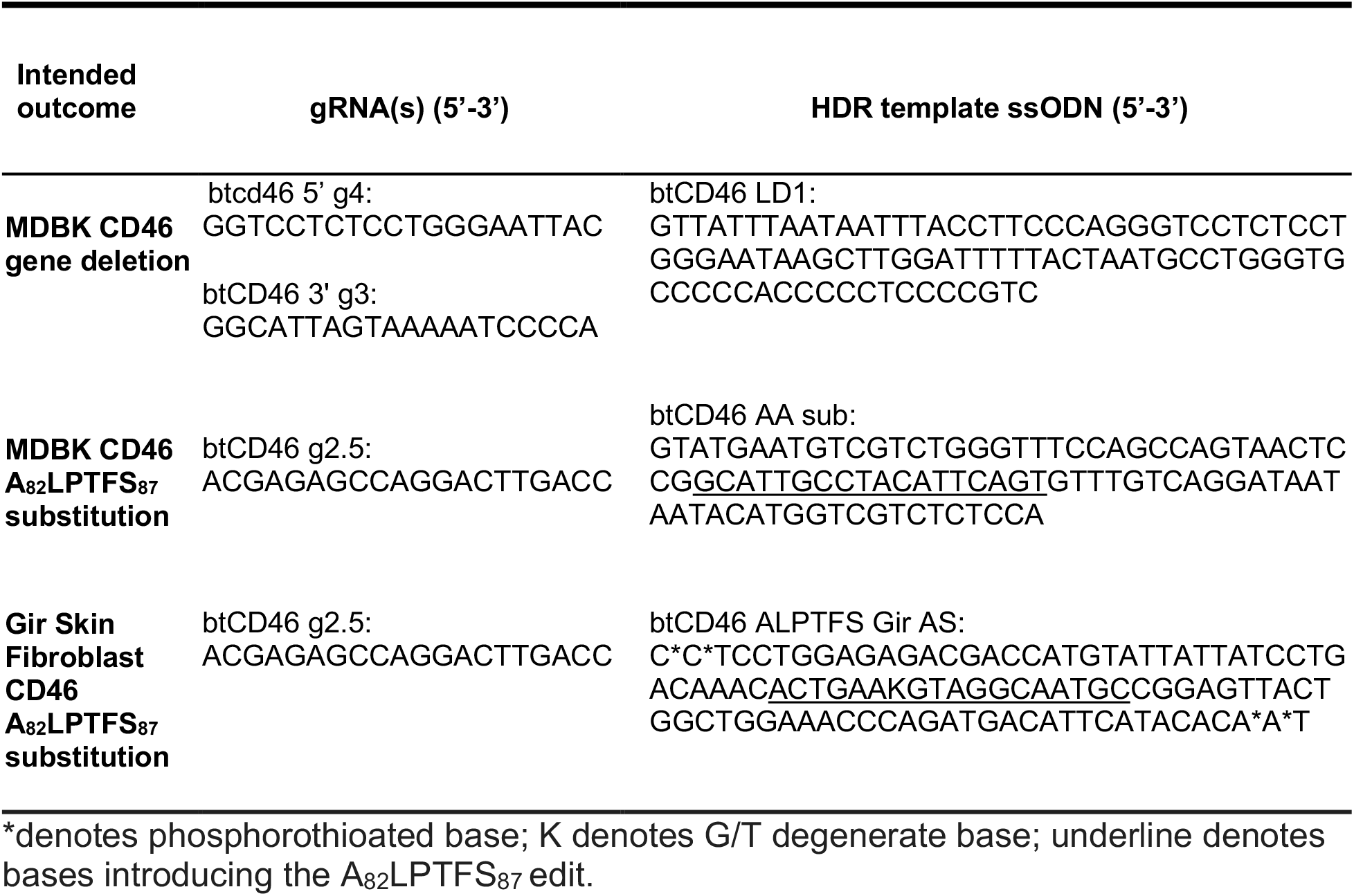
Synthetic guide RNA (gRNA) and single-stranded oligonucleotide donor (ssODN) molecules used in gene-editing.

For the CD46 gene deletion, homology-directed repair (HDR) templates were purchased from Integrated DNA Technologies (IDT; San Jose, CA). gRNA and Cas9 mRNA were synthesized by *in vitro* transcription. pDR274 transcription plasmids containing the gRNA sequences were linearized with DraI (NEB; Ipswich, MA) and amplified using Accustart Taq DNA Polymerase HiFi (Quanta Biosciences, Gaithersburg, MD) using the following primers and cycling program: pDR274 F (5’-TCCGCTCGCACCGCTAGCT-3’) and pDR274 R (5’-AGCACCGACTCGGTGCCAC-3’); 1 cycle (95°C, 2 minutes), 35 cycles of (95°C, 30 s; 48°C, 15 s; 68°C, 15 s), 1 cycle (68°C, 30 s). Once completed, the amplification reactions were treated with RNAsecure (ThermoFisher Scientific; Waltham, MA) following the manufacturer’s recommendations and purified using the QIAquick Gel Extraction Kit (Qiagen; Hilden, Germany). These amplicons were used as a template for transcription using the MEGAshortscript T7 Transcription Kit (ThermoFisher Scientific) and purified with the RNeasy Mini Kit (Qiagen), following manufacturer’s instructions. This RNA transcript constituted the gRNA.

For Cas9 mRNA synthesis, pT3Ts-nCas9n plasmid was linearized with XbaI (NEB), treated with RNAsecure (ThermoFisher Scientific) and purified using the QIAquick Gel Extraction Kit (QIAGEN). *In vitro* Cas9 mRNA transcription was performed using the mMESSAGE mMACHINE T3 Kit (ThermoFisher Scientific) followed by A-tailing using a Poly(A) Tailing Kit (ThermoFisher Scientific). These transcripts were purified with the RNeasy Mini Kit (Qiagen), following manufacturer’s instructions. The RNA transcript constituted the JECas9 mRNA.

For CD46 A_82_LPTFS_87_ substitutions, gRNAs (Alt-R® CRISPR-Cas9 crRNA and Alt-R® CRISPR-Cas9 tracrRNA), HDR templates (single-stranded oligo donor (ssODN)), and Cas9 protein (Cas9 Nuclease V3) were purchased from IDT. The relative position of the gRNA and the ssODN in relation to CD46 protein domains and amino acids sequence are shown in **Fig. 1A and B**. Ribonucleoprotein (RNP) complexes were formed according to the manufacturers recommendations and incubated for 20 minutes immediately prior to transfection.

**Figure 1.**
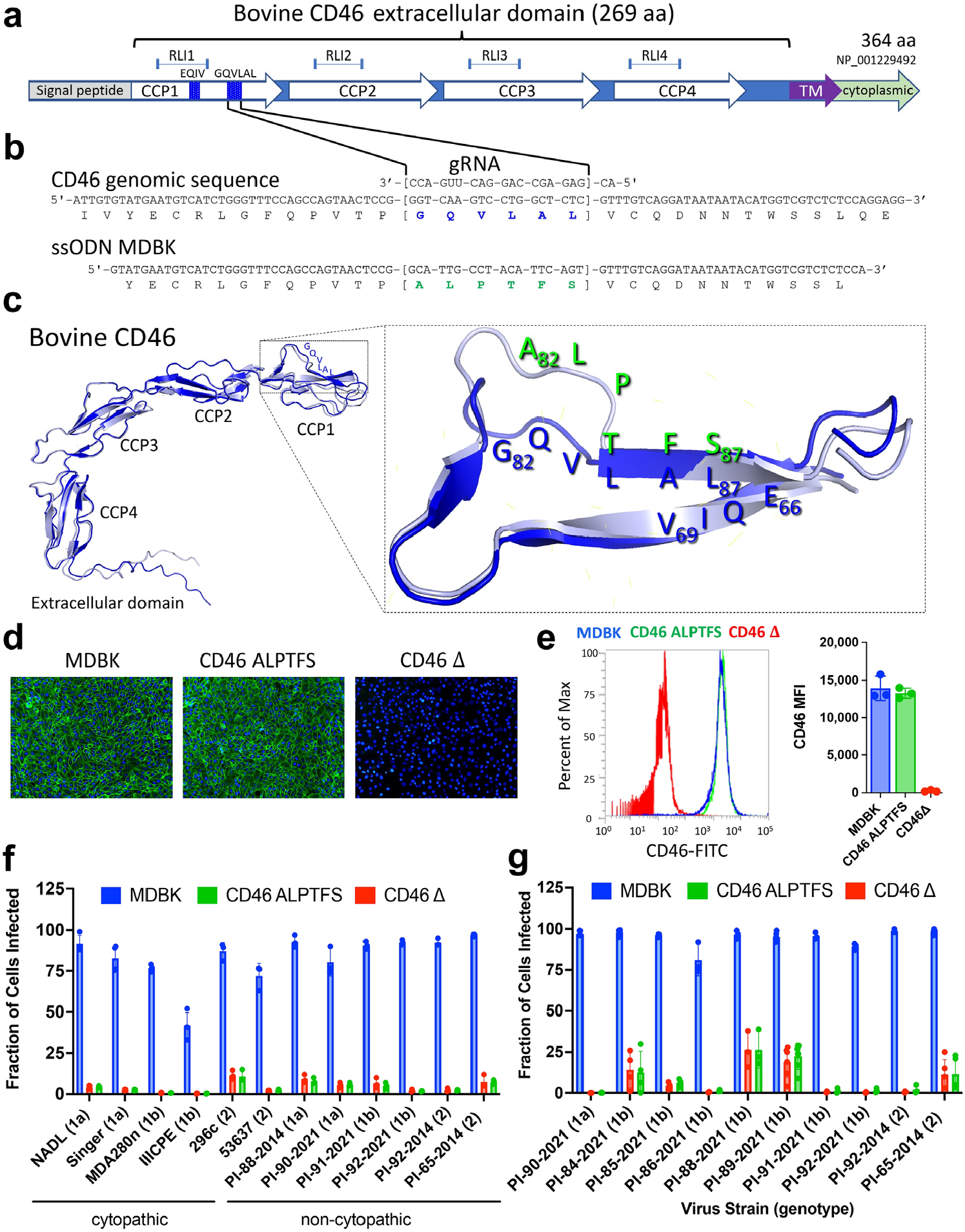
Comparing the CD46 A_82_LPTFS_87_ substitution with CD46 gene deletion in MDBK cells. Panel **a**, physical map of CD46 protein domains. Abbreviations: RLI, receptor-ligand interaction domain; CCP, complement control protein domain; TM, transmembrane domain. Panel **b**, genomic DNA sequence of CD46 in the region of the G_82_QVLAL_87_ motif showing the amino acid translation, the gRNA heteroduplex, and the alignment of the synthetic ssODN used for replacing the G_82_QVLAL_87_ residues with A_82_LPTFS_87_. Abbreviations: gRNA, synthetic guide RNA; ssODN, single stranded oligonucleotide donor template. Panel **c**, ribbon representations of AlphaFold2 predictions of the extracellular domain of wild-type bovine CD46 (blue) aligned with that containing the A_82_LPTFS_87_ substitution (silver). The inset shows a closeup of the BVDV binding platform rotated to visualize the predicted structure differences in residues 82-84. Panel **d**, immunofluorescence staining of CD46 (green) and nuclei (blue; 10x magnification). Panel **e**, flow cytometric quantification of CD46 expression levels (n = 3). Abbreviation: MFI, median fluorescence intensity. Panel **f**, CD46-edited cells were infected with cytopathic or non-cytopathic BVDV isolates at an MOI of 2 and infection efficiency was determined at 20 hpi by flow cytometry using a monoclonal anti-BVDV E2 antibody. Panel **g**, serum from BVDV-PI calves was inoculated on cells. Infection efficiency was quantified at 72 hpi by flow cytometry. In Panels f and g, the results represent the mean ± standard deviation of n ≥ 3 independent experiments.

#### MDBK cell culture and transfection

MDBK cells were maintained at 38.5°C, 5% CO_2_ in DMEM supplemented with 10% irradiated fetal bovine serum, 100 I.U./mL penicillin and streptomycin, and 10mM Hepes. Actively growing cells were prepared for transfection with the Neon Transfection system (Life Technologies; Carlsbad, CA). Briefly, 600,000 cells were resuspended in buffer “R.” For the CD46 gene deletion transfection, 1μg of btCD46 5’ g4 RNA, 1μg of btCD46 3’ g3 RNA, 2μg of JECas9 mRNA, and 0.2nmol of btCD46 LD1 ssODN were added to the cells. For the CD46 A_82_LPTFS_87_ substitution, 120 pMol gRNA:tracrRNA complex, 17.2 μg Alt-R® S.p. HiFi Cas9 Nuclease V3, and 0.2 nmol of the ssODN were added to the cells. The cell suspension was electroporated using the 100μl Neon tip with the following parameters: input voltage: 1600V; pulse width: 20 ms; pulse number: 1. Transfected cells were dispersed into one well of a 6-well plate with 2 ml DMEM media and cultured for 4 days at 30°C.

#### Fibroblast culture and transfection

Primary Gir fibroblast cells were maintained at 37°C, 5% CO_2_ in DMEM supplemented with 20% irradiated fetal bovine serum and 100 I.U./mL penicillin and streptomycin. Actively growing cells were prepared for transfection with the Neon Transfection system (Life Technologies). Briefly, 600,000 cells were resuspended in buffer “R.” For the CD46 A_82_LPTFS_87_ substitution, 65 pmol of gRNA, 78pmol of Alt-R® S.p. HiFi Cas9 Nuclease V3, and 0.11nmol of the ssODN were added to the cells. The cell suspension was electroporated using the 100μl Neon Tip with the following parameters: input voltage: 1800V; pulse width: 20ms; pulse number: 1. Transfected cells were dispersed into one well of a 6-well plate with 2 mL DMEM media and cultured for 3 days at 37°C.

#### Single-cell derived clonal isolation and genotyping

After 4 days at 30°C for the MDBK cells or 3 days at 37°C for the fibroblasts, cells were seeded at low density, subcultured, and screened for homology directed repair by PCR. Screening for homozygous CD46 A_82_LPTFS_87_ in MDBK and fibroblast cells was conducted using AccuStart II GelTrack PCR SuperMix (Quanta Biosciences) and the following primers and cycling program: btCD46 NJ F1 (5’-TTCTCCAACAGGCCAGAAGC -3’) and btCD46 NJ R1 (5’-AGGCAACCAATCGTGACGAA -3’); 1 cycle (95°C, 2 minutes), 35 cycles of (95°C, 20 s; 62°C, 20 s; 72°C, 45 s), 1 cycle (72°C, 5 min). Amplicons were then digested with MspI (NEB) and visualized using agarose gel electrophoresis. Clones homozygous by RFLP were verified by Sanger sequencing (ACGT Inc.; Wheeling, Illinois or Eurofins; Louisville, Kentucky).

Screening for homozygous CD46 gene deletion in MDBK cells was conducted using AccuStart II GelTrack PCR SuperMix (Quanta Biosciences) and the following primers and cycling program: btCD46 5’ F2 (5’-CCAGCCCGCAATGTTTACAC-3’) and btCD46 5’ R2 (5’-TTTGGCCATTGCTCTCCCAA-3’); 1 cycle (95°C, 2 minutes), 35 cycles of (95°C, 20 s; 64°C, 20 s; 72°C, 45 s), 1 cycle (72°C, 5 min). btCD46 5’ F2 (5’-CCAGCCCGCAATGTTTACAC-3’) and btCD46 3’ NJ R1 (5’-TTCACATGCAGACGTGGACT-3’); 1 cycle (95°C, 2 minutes), 35 cycles of (95°C, 20 s; 64°C, 20 s; 72°C, 45 s), 1 cycle (72°C, 5 min). Clones homozygous by junction PCR were verified by Sanger sequencing (ACGT).

### Somatic cell nuclear transfer (SCNT) and reproductive cloning

SCNT was performed by TransOva Genetics (Sioux Center, IA) as previously described^22,23^ on Gir fibroblast clones confirmed homozygous for the A_82_LPTFS_87_ substitution and wild-type unedited controls. Grade 1 embryos containing either the unedited wild-type (n = 8) or CD46 A_82_LPTFS_87_ substitution (n = 8) were implanted into each of 16 recipient cows. Pregnancies were confirmed by ultrasound at 30- and 90-days post-transplantation.

### Fetal tissue collection and primary cell isolation

Fetal tissues were collected at 100-days of gestation from one CD46 A_82_LPTFS_87_ edited fetus and one unedited control fetus and primary cells were isolated from lungs, heart, small intestine, esophagus, kidney, and liver^24–29^. Remaining fetuses were allowed to continue gestation with the goal of reaching full-term.

Primary cells were isolated and maintained in various cell culture mediums. Alveolar cells were maintained in low glucose DMEM (Corning, Corning, NY) supplemented with 20% irradiated fetal bovine serum and 1x antibiotic-antimycotic and cultured on plates coated with 2% gelatin (Sigma, St. Louis, MO). Renal epithelial cells were maintained in DMEM/Hams F-12 50/50 medium (Corning) supplemented with 10% irradiated fetal bovine serum, 1x antibiotic-antimycotic, 5 ug/mL transferrin (Sigma), 25 ng/mL rhEGF (Invitrogen), 0.1 ug/mL hydrocortisone (Sigma) and 10 ug/mL bovine insulin (Sigma). Renal epithelial cells were cultured on plates coated with collagen (Sigma). Small intestine epithelial cells were cultured in the same medium as listed for renal epithelial cells and cultured on plates coated with 0.1% gelatin. Esophageal fibroblasts were cultured in DMEM supplemented with 15% irradiated fetal bovine serum and 1x antibiotic-antimycotic. Epicardial cells were maintained in DMEM + M199 (ThermoFisher Scientific) mixed 1:1 and supplemented with 10% irradiated fetal bovine serum, 1x antibiotic-antimycotic, and 10 uM SB43152 (Sigma). Liver cells (mixed population) were cultured in the same medium as listed for the renal and intestinal epithelial cells and cultured on plates coated with 0.1% gelatin. All cells were tested for BVDV contamination prior to experimentation by RT-qPCR with a BVDV specific primer/probe set ^30^ as described previously^31^.

### Birth of the CD46-edited calf

A CD46 A_82_LPTFS_87_ edited Gir calf was delivered by cesarean section at the calculated time of full gestation (approximately 285 days for Gir cattle). The calf was removed from the recipient cow without nursing and was fed a commercial bovine colostrum replacement solution (Calf’s Choice Total Gold) by bottle. Thereafter, the calf was fed a commercial milk replacement solution formulated for calves. Simultaneously, an age- and sex-matched Holstein dairy calf was purchased from an Iowa dairy farm and fed the same commercial replacement solutions. At one week of age, both calves were moved to the BSL2 animal care facility at the University of Nebraska-Lincoln and housed in the same room.

### Evaluating CD46 DNA sequences of cell lines and animals with 15-fold whole genome sequence (WGS)

DNA was extracted from cell lines or tissues with standard procedures that used RNase/protease digestion, phenol/chloroform extraction, and ethanol precipitation. Purified DNAs were dissolved in a solution of 10 mM TrisCl, 1 mM EDTA (TE, pH 8.0), and stored at 4°C. For WGS, 2 μg of genomic DNA was fragmented and used to make indexed, 500 bp, paired-end libraries. Pooled, indexed libraries were sequenced with massively parallel sequencing machines (either NextSeq500 or NextSeq2000, Illumina Inc.) and the appropriate kits producing 2 x 150 bp paired-end reads. Samples were repeatedly sequenced to a threshold of 40 gigabases surpassing Q20 quality. This approach produced at least 12-fold mapped read coverage (15-fold average) and provided genotype scoring rates and accuracies that exceed 99%^32,33^.

FASTQ files were aggregated by animal and aligned individually to a bovine reference genome (ARS-UCD1.2) with the Burrows-Wheeler Alignment tool (BWA) aln algorithm version 0.7.1219. The files were merged and collated with the bwa sampe command and the resulting sequence alignment map (SAM) files were converted to binary alignment map (BAM) files and sorted with SAMtools version 1.3.120. PCR duplicates were marked in the BAM files with the Genome Analysis Toolkit (GATK) version 3.621. The GATK module RealignerTargetCreator was used to identify regions with small indels, and those regions were realigned with the IndelRealigner module. The BAM files produced at each step were indexed with SAMtools and made available via object storage through a web service interface (Amazon Web Services, Inc. Seattle, WA).

Aligned genomes were viewed with the Integrative Genomics Viewer (IGV) version 2.12.2 by selecting the desired reference genome and loading a session file URL. For example, the TO470 cell line and its unedited and edited reproduction clones were viewed together in IGG by loading session URL: https://s3.us-west-2.amazonaws.com/usmarc.heaton.public/WGS/CellLines/ARS1.2/sessions/ARS1.2_CD46_Ginger4Tracks.xml. The genomic DNA sequences in a 150kb region centered on the *CD46* gene were visually inspected in IGV for differences at the nucleotide level. The edited site encoding CD46 residues 82 to 87 (ARS-UCD1.2; ch16:75,617,415 to ch16:75,617,432) was also inspected. Potential off-target sites were searched for in the ARS-UCD1.2 bovine reference genome with Cas-OFFinder (version 2.4^34^) and the target gRNA sequence: 5’-ACGAGAGCCAGGACTTGACC-3’. Also, *ad hoc* GATK software analysis of WGS was used in an attempt to independently identify any significant differences in the genome sequences between the unedited Gir TO470 cell line, the unedited 100-day fetal calf, the CD46-edited 100-day fetal calf, and the born live CD46-edited fetal calf. Specifically, GATK-identified indels and structural variants were sorted by those that were unique to the live calf compared to the three other samples. Differences were ranked based on size and manually inspected in IGV.

### Immunofluorescence (IF) staining and flow cytometric detection of bovine CD46

Cellular localization of CD46 was visualized by IF staining and quantified by flow cytometry using a polyclonal anti-bovine CD46 antibody. Antibodies that detect bovine CD46 are not commercially available and were thus generated for use in this study. The sequence encoding the extracellular domain of bovine CD46 (amino acids 43-310) was codon optimized and synthesized with an artificial signal peptide which caused the expressed protein to be secreted from cells. The synthetic gene was cloned into the mammalian expression vector pcDNA3.4 and transfected into mammalian cell cultures (Chinese hamster ovary cells, CHO-S). The secreted CD46 peptide was purified from the culture supernatant and injected into two rabbits as an immunogen for primary immunization and successive boosting. Rabbits were boosted three times before terminal blood collection. Antibodies from each rabbit were purified and quantified individually (Genscript BioTech; Piscataway, NJ).

For microscopy, cells were fixed with 4% paraformaldehyde (PFA) in phosphate buffered saline (PBS) for 10 minutes at room temperature. Cells were blocked using 2% bovine serum albumin diluted in phosphate buffered saline (BSA-PBS) for 1 hour at room temperature. The CD46 polyclonal antibody was used at 5 μg /mL diluted in 2% BSA-PBS. Cells were incubated with the CD46 antibody for 1 hour at room temperature. Following three washes with PBS, cells were incubated with a goat anti-rabbit IgG FITC conjugated secondary antibody (Abcam; Cambridge, United Kingdom; catalog no. ab6717) in 2% BSA-PBS and incubated for 1 hour at room temperature in the dark. Following three PBS washes, the nuclei of the cells were stained with a 300 nM solution of DAPI (4′,6-diamidino-2-phenylindole) in PBS for 10 minutes at room temperature. Cells were visualized using an EVOS FL Auto microscope at 10x magnification (Life Technologies).

For flow cytometry, cells were collected using TrypLE (Gibco; Waltham, MA) and centrifuged for 6 minutes at 300 x g. The supernatant was discarded and 5×10^5^ cells were resuspended in 100 μL of 2% BSA-PBS and incubated for 30 minutes on ice. An anti-bovine CD46 polyclonal antibody was added to the tubes at a concentration of 1 μg/mL and incubated for 30 minutes on ice. The cells were washed three times with PBS and resuspended with 2% BSA-PBS containing a goat anti-rabbit IgG FITC conjugated secondary antibody (Abcam catalog no. ab6717) and incubated for 30 minutes on ice in the dark. Cells were washed three times with PBS and analyzed using an Attune NxT Flow Cytometer (ThermoFisher Scientific) using the Attune cytometric software version 5.1.1.

### Virus isolates and serum from calves persistently infected with BVDV (BVDV-PI)

Information on the virus isolates used in this study can be found in **Table S1**.

#### Cytopathic isolates

BVDV-1a stain NADL (ATCC-VR1422) was purchased from ATCC. BVDV strains Singer (BVDV-1a), MDA280n (BVDV-1b), IIICPE (BVDV-1b), 53637 (BVDV-2), and 296c (BVDV-2) were obtained from the National Animal Disease Center (NADC) collection in Ames, Iowa. Cytopathic BVDV strains were propagated in MDBK cells and quantitated in bovine turbinate cells (BT; ATCC CRL 1390). The infective titer was determined in two replicates using an endpoint dilution assay. Viral stocks were stored in 0.5 to 1mL aliquots at -80°C.

#### Serum from BVDV-PI calves

To test non-cytopathic (NCP) field strains, whole blood was collected from cattle persistently infected with BVDV. Serum was separated at 1,650 x g for 15 minutes at 4°C, aliquoted, and stored at -80°C. Virus from each sample was genotyped by PCR amplification and Sanger sequencing of the 5’ untranslated region (UTR) as previously described^35^.

#### Non-cytopathic isolates

NCP BVDV strains were isolated from serum (this study) or plasma^35^ collected from BVDV-PI cattle. Virus was isolated on MDBK cells and low passage (passage 2-4) viral stocks were stored at -80°C. The infective titer of NCP BVDV isolates was estimated by RT-qPCR using log_10_ dilutions of the virus^31^. A standard curve was made by plotting RT-qPCR Ct values against log_10_ dilutions of the NADL virus with a known infectious titer. Linear regression analysis was performed to create a standard for estimating the approximate titer of NCP stocks of virus.

### IF staining and flow cytometric detection of BVDV antigen

For flow cytometric quantification of infected cells or IF visualization, cells were fixed with 4% PFA in PBS, washed, and processed similar to that described above for CD46 detection with the addition of 0.1% saponin (w/v) to all blocking, staining, and wash buffers for cell permeabilization. An anti-BVDV E2 monoclonal antibody (Creative Diagnostics; Shirley, NY, Catalog no. DMAB28412 or VMRD; Pullman, WA, Catalog no. 348) and an anti-mouse CruzFluor™ 488 (CFL 488) conjugated secondary antibody (Santa Cruz Biotechnology; Dallas, TX, catalog no. SC-533653) were used to detect BVDV antigen positive cells.

### BVDV susceptibility testing in CD46-edited MDBK cells

Cells were seeded one day prior to infection in 24-well or 48-well plates in MEM supplemented with 7.5% horse serum (HS; ATCC). Prior to infection, cell culture medium was removed. Then, cells were inoculated with BVDV isolates or serum from BVDV-PI calves as described in the figure legends. Virus adsorption and entry was allowed to proceed for 2 hours at 37°C and unbound virus was removed by washing the cells four times with PBS. Cells were incubated with MEM supplemented with 5% HS for the desired time before infection efficiency was determined by IF staining or flow cytometry as described above.

### Isolation and infection of primary skin fibroblasts

Ear punches were collected from the CD46 A_82_LPTFS_87_ edited calf and the unedited control Holstein calf using the Allflex tissue sampling unit for primary skin fibroblast isolation at TransOva Genetics. Primary skin fibroblasts were grown in Dulbecco’s modified Eagle’s Medium (DMEM) supplemented with 15% irradiated fetal bovine serum and 1x antibiotic-antimycotic. For infection studies, primary fibroblasts were seeded in 24-well plates at a density of 5×10^4^ cells per well in 200 μL DMEM supplemented with 1x antibiotic-antimycotic and 2 mM L-Glutamine, and 5% horse serum (HS; ATCC) and incubated 24 hours at 37°C with 5% CO_2_. The following day, cells were infected with NCP BVDV from the serum of BVDV-PI calves for BVDV genotypes 1a and 1b or with low passage cell culture isolates for BVDV genotype 2. Cell culture medium was removed, and cells were inoculated with serum from BVDV-PI calves diluted 1:1 with DMEM or BVDV-2 isolates at an MOI of 0.1 and incubated for 2 hours at 37°C. Cells were then washed four times with PBS to remove unbound virus and cells were incubated in DMEM supplemented with 5% HS at 37°C for 72 hours. Infected cells were quantified by flow cytometry and visualized by IF as described above.

### PBMC and monocyte isolation and *ex vivo* BVDV challenge

Blood was collected from the CD46 A_82_LPTFS_87_ edited Gir calf and the unedited control Holstein calf via jugular venipuncture into syringes containing EDTA as an anticoagulant. PBMC isolation was accomplished using SepMate tubes according to the manufacturer’s instructions with a few modifications (Stemcell Technologies, Cambridge, MA). Each 9 mL of blood was diluted in 15 mL of room temperature PBS and gently mixed for 15 min on a nutator. Fifteen mL of Ficoll-Plaque Plus at a density of 1.077 g/mL (Cytiva, Marlborough, MA) was added to 50 mL SepMate tubes and the diluted blood was overlaid followed by centrifugation at 1,200 x g for 15 minutes at 21°C with the brake on. The PBMC layer was then collected by pouring the top layer into a fresh 50 mL tube. Approximately 30 mL of PBS was added to each tube, bringing the total volume to 50 mL, and cells were pelleted at 500 x g for 15 minutes at 21°C. Residual red blood cells were lysed using red blood cell lysis buffer (Sigma) according to the manufacturer’s instructions. PBMC were then washed three times by centrifugation (300 x g, 8 min, 21°C) with 50 mL PBS. On the fourth wash, cells were pelleted at 120 x g for 10 min at 21°C to remove platelets from the PBMC preparation. Cell pellets were then resuspended in RPMI plus 1x antibiotic-antimycotic (Gibco) and differential counts were obtained with the Heska Element HT5 (Heska; Loveland, CO).

For monocyte isolation, PBMCs were resuspended at 5×10^5^ monocytes per mL and 700 μL of PBMC suspension containing 3.5×10^5^ monocytes were added to each well in a 48-well plate. Plated cells were incubated at 37°C with 5% CO_2_ for 2 hours. Non-adherent cells (mostly lymphocytes) were aspirated with a pipette and wells were washed vigorously four times with RPMI plus 1x antibiotic-antimycotic to remove all non-adherent cells, leaving behind the monocytes. Two hundred fifty μL of RPMI supplemented with 1x antibiotic-antimycotic and 5% (v/v) heat-inactivated FBS was added to each well and incubated overnight at 37°C with 5% CO_2_.

Non-adherent cells collected from the 48-well plates following monocyte isolation were pelleted at 400 x g for 6 min at 4°C and resuspended in RPMI supplemented with 1x antibiotic-antimycotic and 10% (v/v) heat-inactivated FBS. Cell counts and viability were determined using the Heska Element HT5 and the automated cell counter, Countess II. Cells were then resuspended at 1.25×10^6^ cells per mL and 200 μL of the cell suspension containing 2.5×10^5^ cells was plated in 96-well round bottom plates. Plated cells were incubated overnight at 37°C with 5% CO_2_.

For infection of monocytes, medium was removed from the wells and replaced with 100 μL of RPMI with 10% (v/v) serum collected from persistently infected calves (genotypes 1a and 1b) or BVDV-2 isolate PI-92-2014 at an MOI of 0.01. Monocytes were inoculated for 2 hr at 37°C with 5% CO_2_ with gentle agitation of the plates every 20 min. Two hundred fifty μL of RPMI supplemented with 1x antibiotic-antimycotic and 5% (v/v) heat-inactivated FBS was then added to each well and cells were incubated for 48 hr at 37°C with 5% CO_2_. Input (t=0) samples were also collected and stored at -80°C. Duplicate plates were frozen at 48 hpi and processed for viral RNA detection. Following two freeze thaw cycles to release viral RNA from infected cells, RNA was extracted from clarified supernatants using the Qiagen viral RNA spin columns per the manufacturer’s instructions. Viral RNA was quantified by RT-qPCR with a BVDV specific primer/probe set ^30^ as previously described^31^. Cycle threshold (Ct) values less than 38 were considered positive. Positive, negative, no template, and extraction controls were included on each run. The fold increase in viral RNA compared to the input concentration was determined with the delta Ct method ^36^ and graphed in Prism (v6, GraphPad Software; San Diego, CA).

For infection of lymphocytes, 80 μL of the medium was removed from each well and replaced with 80 μL of various virus isolates at varying MOI (average MOI = 4) so as to add the maximum amount of each virus isolate to the cells. Cells were incubated 24 hours at 37°C with 5% CO_2_ then processed for flow cytometric quantification of BVDV infected cells as described above.

### Natural BVDV exposure challenge study

Serum titers of BVDV neutralizing antibodies were monitored in the CD46 A_82_LPTFS_87_ edited Gir calf and the control Holstein calf, since the colostrum replacement solution contained anti-BVDV antibodies that were passively transferred during their first 24h hours after birth. The natural exposure challenge was initiated only after the BVDV-1 and BVDV-2 serum neutralization antibody titers were negligible (≤ 1:2) in both calves.

For the challenge study, a calf persistently infected with BVDV was purchased from a dairy farm in Iowa. In routine testing, the calf tested positive for BVDV by RT-qPCR on pooled ear notches and by immunohistochemical detection on a fresh ear sample collected 10 days after the initial positive test. To ensure the animal was persistently and not transiently infected, infection status was confirmed after purchase by BVDV RT-qPCR on serum and nasal swab samples collected more than two weeks after the original positive sample. In addition, infectious virus was isolated on MDBK cells from both serum and nasal swab samples using standard protocols (**Fig. S6**). The complete viral sequence was determined by next generation sequencing on the Illumina platform as previously described^35,37^. The virus genotype was determined by sequence alignment and phylogenetic analysis with known reference strains^35^. The calf was approximately 30 days of age and weighed approximately 60 pounds at the time of the challenge study.

The CD46-edited Gir calf and the unedited control Holstein calf were challenged with BVDV by cohabitation with the BVDV-PI calf for 7 days in a BSL2 animal room. Samples collected throughout the study included blood, nasal swabs, and fecal swabs as outlined in **Fig. 4A**. Blood was collected by jugular venipuncture into syringes containing EDTA as an anticoagulant and Sarstedt Monovet tubes (Sarstedt, Inc; Newton, NC) for serum separation. Whole blood samples with EDTA were evaluated using a HT5 veterinary hematology instrument. A 1 mL aliquot of EDTA blood was stored at -80°C for BVDV RT-qPCR. PBMC were isolated from 20 mL EDTA blood using SepMate tubes similar to that described above but with minor modifications. Isolated PBMC were washed once with PBS then counted and differentiated using the Heska instrument. Cells were diluted to 3×10^6^ PBMC per 0.25 mL and stored in four replicate tubes at -80°C for detection of viral RNA by RT-qPCR. Serum was separated from clotted blood by centrifugation at 1660 x g for 15 min at 4°C and aliquoted into tubes and stored at -80°C. Nasal samples were collected by inserting two 6-inch nasal swabs into each nostril approximately 4 inches and gently twisted back and forth for approximately 3 to 5 seconds. One swab from each nostril was placed into duplicate cryovials containing 1mL minimal essential medium (MEM; Gibco) and stored at -80°C. Fecal samples were collected by inserting one swab into the rectum approximately 5 to 7 cm and gently swabbing in a circular motion. The fecal swab was placed into a cryovial containing 1mL MEM and stored at -80°C. BVDV-1 and BVDV-2 virus neutralization assays were completed at the Nebraska Veterinary Diagnostic Center (University of Nebraska-Lincoln) on blinded serum samples.

**Figure 2.**
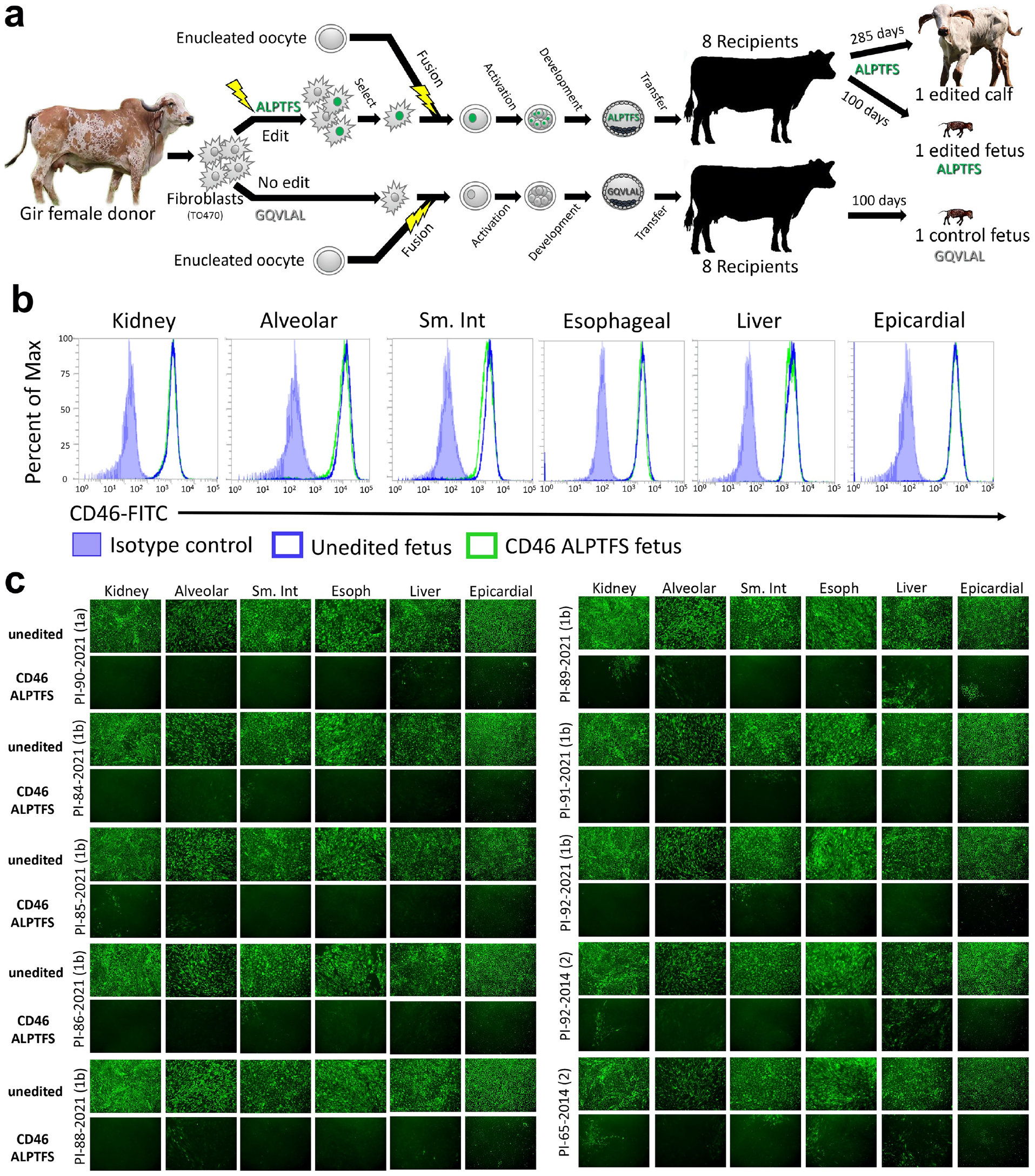
Reproductive cloning and BVDV susceptibility testing of primary cells from 100-day fetal tissues. Panel **a**, schematic representation of reproductive cloning. Primary skin fibroblasts were edited and subsequently fused to enucleated oocytes (somatic cell nuclear transfer) and the resultant embryos implanted into synchronized recipient cows. Panel **b**, flow cytometric quantification of CD46 surface expression. Panel **c**, cells were inoculated with serum (genotypes 1a and 1b) or low passage virus isolates (genotype 2) from BVDV-PI calves. Infection efficiency was determined at 48 hpi using an anti-BVDV monoclonal antibody and FITC labeled secondary antibody. Nuclei were stained with DAPI to ensure images were taken in regions with complete cell monolayers (not shown). Cells imaged at 10x magnification. Abbreviations: Sm. Int, small intestine; Esoph, esophageal.

**Figure 3.**
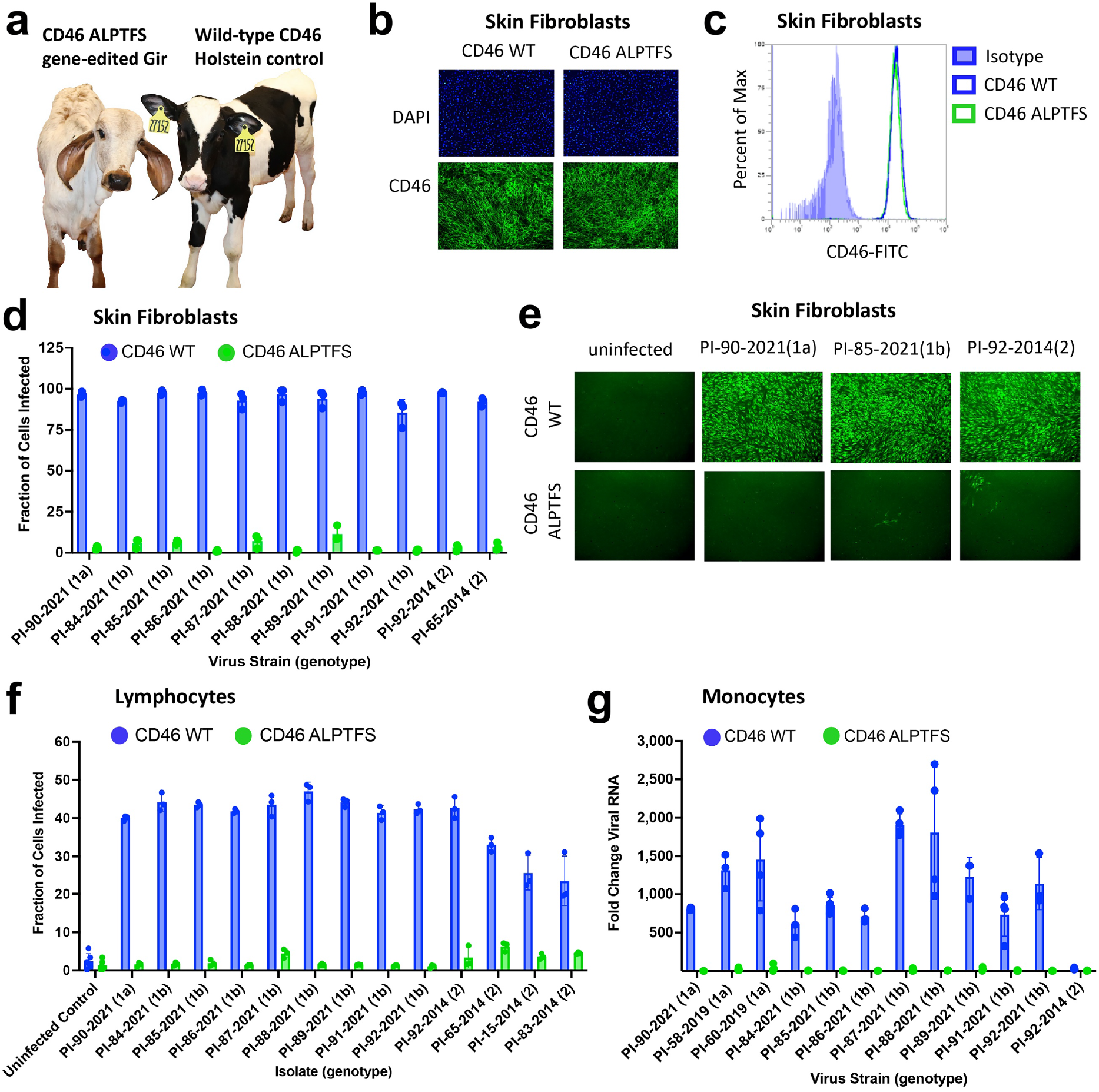
*Ex vivo* challenge of fibroblasts, monocytes, and lymphocytes from CD46 A_82_LPTFS_87_ edited calf. Panel **a**, picture of study participants. Panel **b**, immunofluorescence staining of CD46 (green) and nuclei were stained with DAPI (blue). Panel **c**, flow cytometric quantification of CD46 expression levels. Panel **d**, cells were inoculated with serum from BVDV-PI calves and infection efficiency was quantified at 72 hpi by flow cytometry using an anti-BVDV E2 antibody. Results represent the mean ± standard deviation of n = 3 independent experiments. Panel **e**, BVDV infection was visualized for a representative set of samples from panel d by IF using an anti-BVDV antibody and FITC labeled secondary antibody (10x magnification). Panel **f**, flow cytometric quantification of BVDV infected cells at 24 hpi. Results represent the mean ± SD of n = 3 independent experiments. Panel **g**, monocytes were inoculated with serum from BVDV-PI calves (genotypes 1a and 1b) or a low passage BVDV-2 isolate at an MOI of 0.01 (PI-92-2014). Viral RNA was detected by RT-qPCR at 48 hpi and fold change in viral RNA relative to the input sample (0 hpi) was calculated. Results represent the mean ± SD of n ≥ 3 independent experiments. Abbreviations: WT, wild-type.

**Figure 4.**
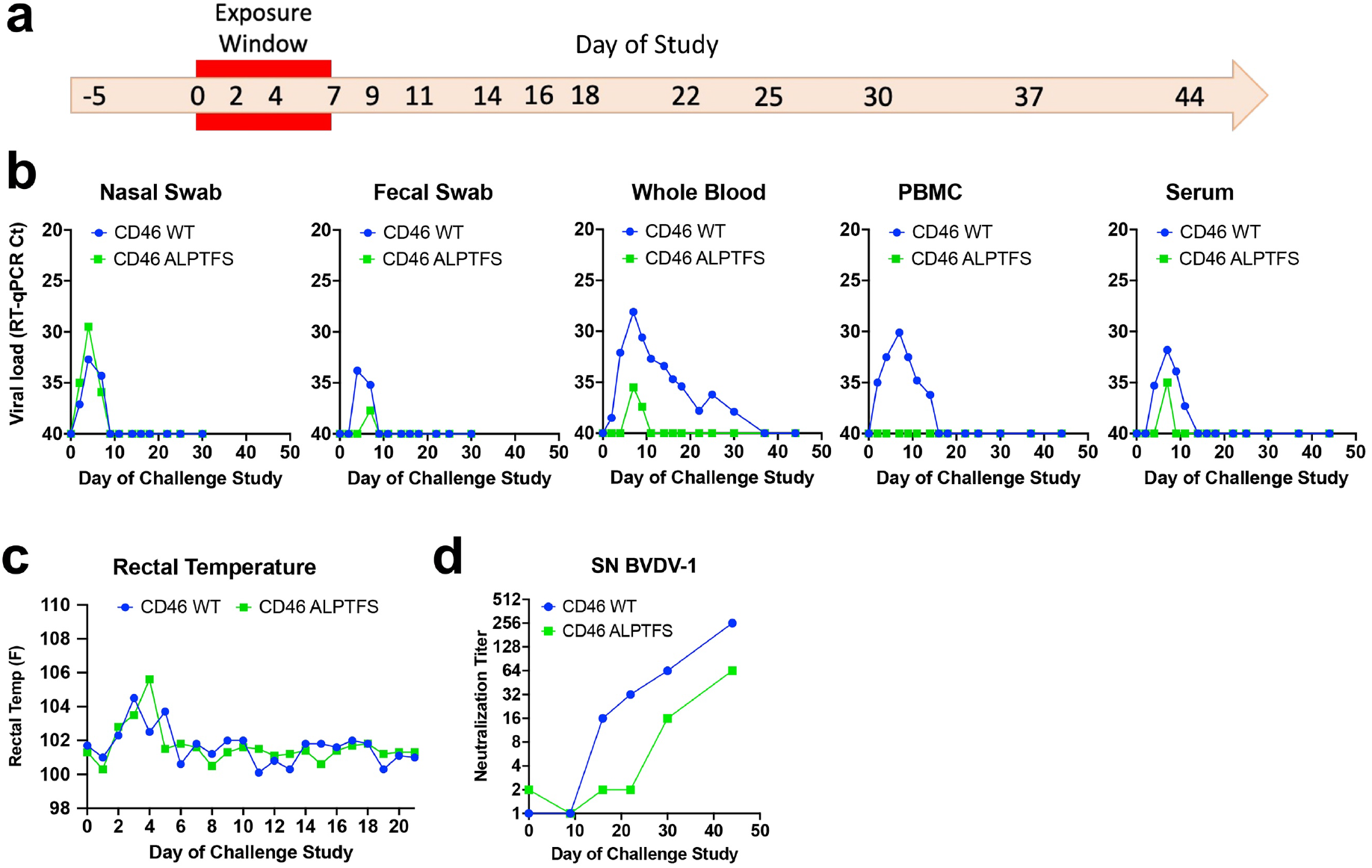
Natural exposure challenge study of CD46 A_82_LPTFS_87_ edited calf. Panel **a**, timeline of challenge study. The CD46-edited Gir calf and the unedited wild-type (WT) control Holstein calf were challenged with BVDV by cohabitation with a BVDV-PI calf for 7 days (red exposure window). Nasal and fecal samples were collected on days -5 to 30. Blood was collected on days -5 to 44. Panel **b**, BVDV RT-qPCR. Samples were extracted in duplicate and run on the same PCR plate. Samples that tested negative for BVDV RNA are plotted having a cycle threshold (Ct) equal to 40. Panel **c**, rectal temperatures taken daily through day 21. Panel **d**, BVDV-1 serum neutralization (SN) titers.

In addition to sample collections by researchers, an animal health technician spent approximately 30 minutes in the room daily to provide a clinical assessment of each calf’s health. A score ranging from 0 to 16 was determined for each animal based on attitude, presence of cough, nasal discharge, eye discharge, fecal consistency, and rectal temperature according to a modified Wisconsin Health Scoring System^38^ (**Table S2**).

### Statistical Analyses

Statistical analyses were performed using GraphPad Prism version 9.4.0 (GraphPad Software Inc.). Comparisons of means plus or minus the standard error (± SE) were analyzed using two-way analysis of variance (ANOVA) followed by Tukey’s (three or more columns) or Sidak’s (two columns) multiple comparisons test. For all comparisons, a *P* value less than 0.05 was considered significant.

## Results

### The effect of the CD46 A_82_LPTFS_87_ substitution in MDBK cells

CRISPR/Cas9 gene-editing by homology-directed repair was used to generate MDBK cells with the A_83_LPTFS_88_ substitution in the BVDV binding domain of CD46 (**Fig. 1A and 1B**). Alignment of folded bovine CD46 protein structures containing the wild-type G_82_QVLAL_87_ or the A_82_LPTFS_87_ substitution showed a predicted structural change limited to the A_82_L_83_P_84_ site (**Fig. 1C**), however CD46 surface expression remained normal in edited cells (**Fig. 1D and 1E**). For comparison, MDBK cells were also generated with a 42-kb deletion encompassing the entire *CD46* gene (CD46Δ). WGS analysis showed the A_82_LPTFS_87_ and the CD46Δ edits were accurate (**Fig. S1**), and off-target site modifications were not found in 125 potential genome sites when allowing for up to 4 bp mismatches with the gRNA. Both the CD46Δ and CD46 A_82_LPTFS_87_ cells showed a dramatic reduction in susceptibility to BVDV compared to the parent MDBK cell line, regardless of the BVDV strain type (**Fig. 1F, 1G, and S2**). For example, infection efficiency was reduced by 95% on average (range 87-99%) across all virus isolates in the CD46-edited cells compared to MDBK at 20 hpi (**Fig. 1F**) and by 92% (range 73-99%) at 72 hpi (**Fig. 1G**). Conversely, infection efficiency was equivalent between CD46Δ and CD46 A_82_LPTFS_87_ edited cells for all but one BVDV strain tested (PI-85-2022, *P* = 0.03, **Fig. 1G**). Together, these data suggest that bovine CD46 with the A_82_LPTFS_87_ substitution may reduce BVDV infection similar to that of a CD46 gene deletion, while allowing CD46 to be expressed normally at the cell surface.

### Production of gene-edited CD46 A_82_LPTFS_87_ cloned calves

To test the impact of the CD46 A_82_LPTFS_87_ edit *in vivo*, the same strategy was used to make the edit in a primary Gir fibroblast cell line (TO470) for reproductive cloning (**Fig. 2A**). Three of eight embryos of each type (CD46-edited and unedited control) had established successful pregnancies 30-days after implantation. However, at 90-days, two of the unedited fetuses had been lost and resorbed. At 100 days gestation, one unedited and one CD46 A_82_LPTFS_87_ edited fetus were collected for primary cell isolation from lung, heart, small intestine, esophagus, kidney, and liver tissue. The two remaining CD46 A_82_LPTFS_87_ edited fetuses continued gestating to test for normal calf development. One edited CD46 A_82_LPTFS_87_ fetus was spontaneously aborted at 212 days gestation but had no obvious physical defects (**Fig. S3**). The last remaining CD46 A_82_LPTFS_87_ edited Gir calf was delivered by cesarean section at full term (285 days) and was born healthy on July 19, 2021 (**Fig. S4**). This calf has continued to thrive since and provided the first evidence that a CD46 A_82_LPTFS_87_ substitution may not negatively impact pre- or postnatal development through at least 16 months of age.

### Off-target editing not detected in CD46-edited calves

Analysis of WGS did not reveal any off-target edits in the edited CD46 A_83_LPTFS_88_ 100-day fetus or live-born calf. Except for the intended A_83_LPTFS_88_ substitutions, the 42 kb region spanning *CD46* was homozygous and identical in the donor Gir fibroblast line, the unedited fetus, the edited fetus, and the live-born calf (**Fig. S5**). The first heterozygous SNPs upstream and downstream of *CD46* are at 42 and 48 kb, respectively (chr16:75,663,583 and chr16:75,572,857) and are also identical in all four samples. No off-target sites were found matching the gRNA in the bovine reference genome (ARS-UCD1.2) when zero, one, or two mismatches with zero bulge size was allowed. When three or four mismatches were allowed, seven and 118 potential genomic sites were identified, respectively. Manual inspection of these 125 sites with IGV showed no genomic sequence differences between the parental Gir fibroblast, the unedited fetus, the CD46-edited fetus, and the live-born calf. Also, systematic GATK software analysis of WGS data did not detect any significant off-target insertions or deletions in the genome sequences that were unique to the live-born edited calf. Thus, phenotypic differences observed in BVDV susceptibility between unedited and edited calves were not readily attributable to off-target site modifications.

### Reduced BVDV susceptibility in CD46-edited fetal tissues

The CD46 A_83_LPTFS_88_ substitution had a significant impact on reducing BVDV susceptibility in primary cells from all tissues from the 100-day fetus. The CD46 A_83_LPTFS_88_ substitution did not appreciably alter the surface expression levels of CD46 when measured in primary cultures by flow cytometry (**Fig. 2B**). Yet, the primary cells from all tissue types with the A_83_LPTFS_88_ substitution had a significant reduction in BVDV infected cells (as visualized by IF) compared to cells from the unedited clone, regardless of BVDV strain type (**Fig. 2C**). Based on these results, multiple tissues/organs in a CD46 A_83_LPTFS_88_ edited calf were expected to have reduced BVDV susceptibility.

### *Ex vivo* challenge of cells from the CD46-edited Gir calf

Three available and relevant cell types were collected from the live-born CD46-edited calf for *ex vivo* challenge: skin fibroblasts, peripheral blood lymphocytes, and monocytes. For comparison, a healthy one-day old Holstein calf was purchased and housed with the CD46-edited Gir calf to serve as an unedited sex- and age-matched control, since no unedited Gir calves survived to term (**Fig. 3A**). CD46 localization (**Fig. 3B**) and expression (**Fig. 3C**) was unaltered in primary skin fibroblasts from the CD46-edited calf, yet infection efficiency was reduced by 96% on average (range 88-99%) as quantified by flow cytometry (**Fig. 3D**) and visualized by IF (**Fig. 3E**). Peripheral blood cells can also be infected by BVDV and these cells are thought to disseminate virus throughout the host. Lymphocytes from the CD46-edited calf had on average a 93% reduction (range 81-98%) in BVDV susceptibility as quantified by flow cytometry (**Fig. 3F**). Similarly, monocytes from the CD46-edited calf had a dramatic reduction in susceptibility to BVDV as quantified by RT-qPCR, with an average 163-fold reduction (range 7 to 446-fold) in viral RNA accumulation (**Fig. 3G**). Thus, each of the three cell types tested from the CD46-edited calf displayed a significant reduction in BVDV susceptibility *ex vivo* and suggested that the edited calf might resist infection in a natural exposure challenge experiment.

### Natural challenge of a CD46-edited Gir calf by exposure to a BVDV-PI Calf

At 10 months of age, the CD46 A_83_LPTFS_88_ edited Gir calf and the wild-type CD46 control Holstein were co-housed with a week-old Holstein calf naturally and persistently infected with BVDV (genotype 1b, **Fig. S6**). Both calves received significant BVDV exposure from the BVDV-PI calf during the seven days of exposure as measured by virus detection in nasal and fecal swabs (**Fig. 4A and 4B**). Both calves also developed a fever (**Fig. 4C**) and had changes in blood cell counts (**Table S4**), but only the wild-type CD46 calf displayed other signs of infection, including a cough, rhinitis, and redness and chafing around the nostrils (**Table S3**). Concurrent with clinical signs of infection, BVDV viremia, as measured by RT-qPCR of whole blood, was detected in the wild-type calf and lasted 28 days. Conversely, BVDV RNA was only detected in the whole blood of the CD46-edited calf for three days, and the peak viral RNA load was reduced approximately 2-log (**Fig. 4B**). Furthermore, PBMC isolated from the CD46-edited calf remained uninfected throughout the challenge whereas PBMC from the wild-type CD46 calf were BVDV RNA positive for 12 days. While low levels of cell-free BVDV RNA could be detected in the serum of both calves (**Fig. 4B**), infectious virus could not be isolated from the serum when inoculated on MDBK cells and blindly passaged three times. In contrast, replication-competent virus was easily isolated from the serum of the BVDV-PI calf (**Fig. S6**). Both calves generated neutralizing antibodies to BVDV (**Fig. 4D**), indicating that the CD46 A_83_LPTFS_88_ edited Gir calf’s immune system was competent to respond to the viral challenge. Together, these results suggest that in the face of overwhelming natural BVDV exposure, the CD46 A_83_LPTFS_88_ edited Gir calf displayed significantly reduced susceptibility to BVDV which resulted in no observable adverse health effects on the animal.

## Discussion

The present report describes the use of CRISPR/Cas9 technology to make an A_82_LPTFS_87_ substitution in the BVDV binding domain of CD46. In stepwise experiments, we showed this edit dramatically reduced BVDV susceptibility in MDBK cells, primary cells from multiple organs obtained from a CD46-edited fetus, fibroblast and immune cell populations from a live CD46-edited calf, and in a natural exposure challenge study with the same edited calf. Together, these results provide proof-of-concept for using intentional genome alterations in CD46 to reduce the burden of BVDV-associated diseases in cattle. This represents the first demonstration of editing a bovine gene to reduce susceptibility to a viral pathogen.

The substitution of six amino acids in CD46 is unique compared to previous reports of gene-edited cattle with reduced susceptibility to bacterial pathogens. For example, exogenous whole gene-knockin strategies have been used in cattle to reduce mastitis (*Staphylococcus aureus*)^39,40^ and bovine tuberculosis (*Mycoplasma bovis*)^41–43^. In another example, a single amino acid was changed in bovine CD18 (integrin beta 2), causing the normally retained signal peptide to be cleaved^44^. Although leukocytes isolated from a CD18-edited fetus were resistant to *Manheimia haemolytica* leukotoxin when challenged *ex vivo*, live calves were not reported. Here, homology directed repair was used to make a precise 18-nt replacement in the endogenous *CD46* gene. Subsequently, somatic cell nuclear transfer was used to produce an edited calf in a single generation without introducing any off-target modifications. Importantly, the edit did not disrupt the predicted tertiary structure of the protein or its expression levels, and the live CD46-edited calf has no obvious adverse effects from the on-target edit in the first 16 months after birth.

This study had four important limitations. First, only one live CD46-edited animal could be tested for altered BVDV susceptibility. Second, a live unedited wild-type CD46 clone was not available to serve as a matched control. Third, a breed-matched control was not available since Gir are uncommon in the U.S. Although differences in breed-associated BVDV susceptibility have not been reported, ideally, comparisons would be made between clones that were identical in genomic sequence apart from the six amino acid substitution in CD46. Fourth, our *in vivo* challenge study design did not allow direct determination of viral entry and replication in various tissues of the edited calf, since it would have required sacrificing the only available CD46-edited animal. Although nasal and fecal swab samples were collected to detect potential amplification of virus in the upper respiratory and gastrointestinal tracts, samples from both the CD46-edited and control calves were only positive for BVDV on the days they were co-housed with the BVDV-PI calf. Thus, these positive samples could be from environmental exposure and not active viral replication. As such, questions remain about the susceptibility of various tissues *in vivo*. Future studies should include additional animals from the most common beef and dairy breeds to help address these limitations.

Despite the limitations, the concordance of results between the *in vitro* and *ex vivo* BVDV susceptibility studies with the *in vivo* challenge study suggest that this edit could be useful for reducing BVDV susceptibility in cattle. Fetal kidney, lung, small intestine, esophagus, liver, and heart cells all had significantly reduced susceptibility to BVDV when infected *ex vivo*. Consistent with this result, primary skin fibroblasts, lymphocytes, and monocytes from the live CD46-edited calf had dramatically reduced BVDV susceptibility. Of note, the edit reduced susceptibility to all BVDV isolates tested, including cytopathic and non-cytopathic isolates belonging to the two genotypes of BVDV that infect cattle globally. This component of our study was the first to measure CD46-dependence in diverse cell types and provides new insights into CD46 receptor usage. Although the *in vivo* challenge study design did not directly determine viral replication in individual organs and tissues of the edited calf, the strong reduction in both the duration and peak viral RNA load in the blood of the CD46-edited calf was consistent with reduced viral replication in tissues *in vivo*. This coincides with PBMC results from the CD46-edited calf remaining uninfected during the entire challenge study. Thus, although questions remain about viral replication *in vivo*, our data suggests a profound decrease in viral susceptibility that limits the amount of virus in the blood. Although untested, we hypothesize that this reduced viremia would limit the dissemination of virus to other target organs.

Reducing viremia in acute infections may significantly impact BVDV control efforts. The most devastating outcome of transient infection in pregnant animals is viremia leading to transplacental infection of the developing fetus. Fetal infections can result in abortion, congenital malformations, or worse, the birth of BVDV-PI calves. These calves have lifelong viremia and continuously shed virus in all bodily secretions, making them the most important source of virus spread in the population and the target of national eradication programs^45^. In addition, the immunosuppressive effects of BVDV during an acute infection increase the incidence of secondary infection and disease^46^. For example, BVDV is an important cofactor of bovine respiratory disease complex (BRDC),^47,48^ one of the most economically important infectious diseases affecting the cattle industry. Therefore, reducing transplacental BVDV infections could significantly improve animal health and welfare, reduce the burden of BVDV infections on the industry, and provide a significant opportunity to reduce antibiotic use in agriculture. Accordingly, it will be important for future studies to determine whether CD46-edited dams are able to protect their developing fetuses from transplacental BVDV infection.

## Conclusion

Stepwise experiments showed that substituting A_82_LPTFS_87_ in CD46 dramatically reduced BVDV susceptibility *in vitro, ex vivo*, and in a natural challenge study with one edited calf. The results provide proof-of-concept for using intentional genome alterations in CD46 to reduce the burden of BVDV-associated diseases in cattle. However, determining the ability of CD46-edited animals to withstand BVDV viral challenges *in vivo* will require experimental replication in other breeds and with more animals.

## Data and material availability

Raw WGS files (fastq) for the MDBK and TO470 cell lines and their edited clones are available in the NCBI SRA under accession number SRR21834845 - 50. The sequence data has also been deposited with links to BioProject accession number PRJNA887820 (BioSamples SAMN31186839 - 44) in the NCBI BioProject database.

In addition, IGV access to the aligned sequences (bam files and IGV session files) are available:

MDBK: https://s3.us-west2.amazonaws.com/usmarc.heaton.public/WGS/CellLines/ARS1.2/bams/LIB14394_Bovine_MDBKcells.Bt_ARS-UCD1.2.realigned.bam

MDBK CD46 gene deletion: https://s3.us-west-2.amazonaws.com/usmarc.heaton.public/WGS/CellLines/ARS1.2/bams/LIB102516_BovMDBK_CD46KO3.Bt_ARS-UCD1.2.realigned.bam

MDBK CD46 ALPTFS substitution:https://s3.us-west-2.amazonaws.com/usmarc.heaton.public/WGS/CellLines/ARS1.2/bams/LIB109073_BovMDBK_CD46Por6AA_296.Bt_ARS-UCD1.2.realigned.bam

MDBK IGV Session URL: https://s3.us-west-2.amazonaws.com/usmarc.heaton.public/WGS/CellLines/ARS1.2/sessions/ARS12._MDBK_CD46_3tracks.xml

Gir cell line (TO470): https://s3.us-west-2.amazonaws.com/usmarc.heaton.public/WGS/CellLines/ARS1.2/bams/LIB108537_TO470CellLine.Bt_ARS-UCD1.2.realigned.bam

Unedited fetus:https://s3.us-west-2.amazonaws.com/usmarc.heaton.public/WGS/CellLines/ARS1.2/bams/LIB113017_TO470FetalCalfSkin.Bt_ARS-UCD1.2.realigned.bam

CD46 ALPTFS edited fetus: https://s3.us-west-2.amazonaws.com/usmarc.heaton.public/WGS/CellLines/ARS1.2/bams/LIB113018_TO470_CD46_6AAFetalCalfSkin.Bt_ARS-UCD1.2.realigned.bam

CD46 ALPTFS edited live calf: https://s3.us-west-2.amazonaws.com/usmarc.heaton.public/WGS/CellLines/ARS1.2/bams/LIB110781_Ginger.Bt_ARS-UCD1.2.realigned.bam

Gir clone IGV Session URL: https://s3.us-west-2.amazonaws.com/usmarc.heaton.public/WGS/CellLines/ARS1.2/sessions/ARS1.2_CD46_Ginger4Tracks.xml

Biological materials will be made available upon request. However, primary cells from fetal tissues will not be made freely available due to very limited quantity.

## Acknowledgments

We thank Susan Hauver and the USMARC Core Facility for technical support, the University of Nebraska animal care staff, and Janel Nierman for secretarial and administrative support. Mention of trade names or commercial products in this publication is solely for the purpose of providing specific information and does not imply recommendation or endorsement by the U.S. Department of Agriculture. The USDA is an equal opportunity provider and employer.

## Author contributions

Conceptualization, A.M.W., M.P.H., T.S.S.; Data Curation, A.M.W., M.P.H., T.S.K; Formal Analysis, A.M.W., D.A.W., D.F.C., L.S., M.P.H., T.S.K.; Funding Acquisition, A.M.W., M.P.H, T.S.S.; Investigation, A.M.W., B.L.V, D.A.W., E.E.J., L.S., M.P.H.; Methodology, A.M.W., B.L.V., D.A.W., D.F.C., G.P.H., L.S., M.P.H., T.S.K.; Project Administration, A.M.W., D.F.C., M.P.H., S.L., T.S.S.; Resources, B.L.V., E.E.J., G.P.H., T.S.K.; Software, G.P.H., T.S.K; Supervision, A.M.W., B.L.V., D.F.C., M.P.H., T.S.S.; Validation, A.M.W., D.F.C., M.P.H.; Visualization, A.M.W., M.P.H.; Writing – Original Draft Preparation, A.M.W., M.P.H; Writing – Review & Editing, all authors reviewed and approved the final draft.

## References

1. Baker, J. C. The Clinical Manifestations of Bovine Viral Diarrhea Infection. Veterinary Clinics of North America: Food Animal Practice vol. 11 425–445 Preprint at https://doi.org/10.1016/s0749-0720(15)30460-6 (1995).

2. Bolin, S. R. Bovine Viral Diarrhea Virus in Mixed Infections. Polymicrobial Diseases 31–50 Preprint at https://doi.org/10.1128/9781555817947.ch3 (2014).

3. Bruschke, C. J., Weerdmeester, K., Van Oirschot, J. T. & Van Rijn, P. A. Distribution of bovine virus diarrhoea virus in tissues and white blood cells of cattle during acute infection. Vet. Microbiol. 64, 23–32 (1998).

4. Liess, B., Moennig, V., Pohlenz, J. & Trautwein, G. Ruminant Pestivirus Infections: Virology, Pathogenesis, and Perspectives of Prophylaxis. (Springer Science & Business Media, 2012).

5. McClurkin, A. W. et al. Production of cattle immunotolerant to bovine viral diarrhea virus. Can. J. Comp. Med. 48, 156–161 (1984).

6. Ridpath, J. F. Immunology of BVDV vaccines. Biologicals vol. 41 14–19 Preprint at https://doi.org/10.1016/j.biologicals.2012.07.003 (2013).

7. Burkard, C. et al. Precision engineering for PRRSV resistance in pigs: Macrophages from genome edited pigs lacking CD163 SRCR5 domain are fully resistant to both PRRSV genotypes while maintaining biological function. PLOS Pathogens vol. 13 e1006206 Preprint at https://doi.org/10.1371/journal.ppat.1006206 (2017).

8. Whitworth, K. M. et al. Gene-edited pigs are protected from porcine reproductive and respiratory syndrome virus. Nat. Biotechnol. 34, 20–22 (2016).

9. Wells, K. D. et al. Replacement of Porcine CD163 Scavenger Receptor Cysteine-Rich Domain 5 with a CD163-Like Homolog Confers Resistance of Pigs to Genotype 1 but Not Genotype 2 Porcine Reproductive and Respiratory Syndrome Virus. Journal of Virology vol. 91 Preprint at https://doi.org/10.1128/jvi.01521-16 (2017).

10. Krey, T. et al. Function of bovine CD46 as a cellular receptor for bovine viral diarrhea virus is determined by complement control protein 1. J. Virol. 80, 3912–3922 (2006).

11. Chen, H.-W. et al. Viral Traits and Cellular Knock-Out Genotype Affect Dependence of BVDV on Bovine CD46. Pathogens 10, (2021).

12. Szillat, K. P., Koethe, S., Wernike, K., Höper, D. & Beer, M. A CRISPR/Cas9 Generated Bovine CD46-knockout Cell Line—A Tool to Elucidate the Adaptability of Bovine Viral Diarrhea Viruses (BVDV). Viruses vol. 12 859 Preprint at https://doi.org/10.3390/v12080859 (2020).

13. Leveringhaus, E., Cagatay, G. N., Hardt, J., Becher, P. & Postel, A. Different impact of bovine complement regulatory protein 46 (CD46) as a cellular receptor for members of the species and. Emerg. Microbes Infect. 11, 60–72 (2022).

14. Sa-Carvalho, D. et al. Tissue culture adaptation of foot-and-mouth disease virus selects viruses that bind to heparin and are attenuated in cattle. J. Virol. 71, 5115–5123 (1997).

15. Cagno, Cagno, Tseligka, Jones & Tapparel. Heparan Sulfate Proteoglycans and Viral Attachment: True Receptors or Adaptation Bias? Viruses vol. 11 596 Preprint at https://doi.org/10.3390/v11070596 (2019).

16. Liszewski, M. K. & Atkinson, J. P. Membrane cofactor protein (MCP; CD46): deficiency states and pathogen connections. Curr. Opin. Immunol. 72, 126–134 (2021).

17. Liszewski, M. K. & Kemper, C. Complement in Motion: The Evolution of CD46 from a Complement Regulator to an Orchestrator of Normal Cell Physiology. J. Immunol. 203, 3–5 (2019).

18. Yamamoto, H., Fara, A. F., Dasgupta, P. & Kemper, C. CD46: The ‘multitasker’ of complement proteins. The International Journal of Biochemistry & Cell Biology vol. 45 2808–2820 Preprint at https://doi.org/10.1016/j.biocel.2013.09.016 (2013).

19. Jumper, J. et al. Highly accurate protein structure prediction with AlphaFold. Nature 596, 583–589 (2021).

20. Varadi, M. et al. AlphaFold Protein Structure Database: massively expanding the structural coverage of protein-sequence space with high-accuracy models. Nucleic Acids Res. 50, D439–D444 (2022).

21. Park, J., Bae, S. & Kim, J.-S. Cas-Designer: a web-based tool for choice of CRISPR-Cas9 target sites. Bioinformatics 31, 4014–4016 (2015).

22. Kasinathan, P. et al. Effect of fibroblast donor cell age and cell cycle on development of bovine nuclear transfer embryos in vitro. Biol. Reprod. 64, 1487–1493 (2001).

23. Kuroiwa, Y. et al. Cloned transchromosomic calves producing human immunoglobulin. Nat. Biotechnol. 20, 889–894 (2002).

24. McClenahan, D. et al. Effects of lipopolysaccharide and Mannheimia haemolytica leukotoxin on bovine lung microvascular endothelial cells and alveolar epithelial cells. Clin. Vaccine Immunol. 15, 338–347 (2008).

25. Dronkers, E., Moerkamp, A. T., van Herwaarden, T., Goumans, M.-J. & Smits, A. M. The Isolation and Culture of Primary Epicardial Cells Derived from Human Adult and Fetal Heart Specimens. J. Vis. Exp. (2018) doi:10.3791/57370.

26. Ding, W., Yousefi, K. & Shehadeh, L. A. Isolation, Characterization, And High Throughput Extracellular Flux Analysis of Mouse Primary Renal Tubular Epithelial Cells. J. Vis. Exp. (2018) doi:10.3791/57718.

27. Katwal, P., Thomas, M., Uprety, T., Hildreth, M. B. & Kaushik, R. S. Development and biochemical and immunological characterization of early passage and immortalized bovine intestinal epithelial cell lines from the ileum of a young calf. Cytotechnology 71, 127–148 (2019).

28. Spotorno, V. G., Hidalgo, A., Barbich, M., Lorenti, A. & Zabal, O. Culture of bovine hepatocytes: a non-perfusion technique for cell isolation. Cytotechnology 51, 51–56 (2006).

29. Ian Freshney, R. Culture of Animal Cells. (Wiley-Liss, 1993).

30. Mahlum, C. E. et al. Detection of bovine viral diarrhea virus by TaqMan reverse transcription polymerase chain reaction. J. Vet. Diagn. Invest. 14, 120–125 (2002).

31. Workman, A. M. et al. Evaluating Large Spontaneous Deletions in a Bovine Cell Line Selected for Bovine Viral Diarrhea Virus Resistance. Viruses 13, (2021).

32. Kalbfleisch, T. & Heaton, M. P. Mapping whole genome shotgun sequence and variant calling in mammalian species without their reference genomes. F1000Res. 2, 244 (2013).

33. Heaton, M. P. et al. Using diverse U.S. beef cattle genomes to identify missense mutations in EPAS1, a gene associated with pulmonary hypertension. F1000Research vol. 5 2003 Preprint at https://doi.org/10.12688/f1000research.9254.2 (2016).

34. Bae, S., Park, J. & Kim, J.-S. Cas-OFFinder: a fast and versatile algorithm that searches for potential off-target sites of Cas9 RNA-guided endonucleases. Bioinformatics 30, 1473–1475 (2014).

35. Workman, A. M. et al. Resolving Bovine viral diarrhea virus subtypes from persistently infected U.S. beef calves with complete genome sequence. J. Vet. Diagn. Invest. 28, 519–528 (2016).

36. Livak, K. J. & Schmittgen, T. D. Analysis of Relative Gene Expression Data Using Real-Time Quantitative PCR and the 2−ΔΔCT Method. Methods vol. 25 402–408 Preprint at https://doi.org/10.1006/meth.2001.1262 (2001).

37. Workman, A. M., Clawson, M. L., Heaton, M. P. & Dickey, A. M. First Complete Genome Sequence of a Genotype A2, Subgroup 4 Small Ruminant Lentivirus. Microbiol Resour Announc 7, (2018).

38. McGuirk, S. M. Disease Management of Dairy Calves and Heifers. Veterinary Clinics of North America: Food Animal Practice vol. 24 139–153 Preprint at https://doi.org/10.1016/j.cvfa.2007.10.003 (2008).

39. Liu, X. et al. Zinc-finger nickase-mediated insertion of the lysostaphin gene into the beta-casein locus in cloned cows. Nat. Commun. 4, 2565 (2013).

40. Liu, X. et al. Generation of mastitis resistance in cows by targeting human lysozyme gene to β-casein locus using zinc-finger nucleases. Proc. Biol. Sci. 281, 20133368 (2014).

41. Wu, H. et al. TALE nickase-mediated SP110 knockin endows cattle with increased resistance to tuberculosis. Proc. Natl. Acad. Sci. U. S. A. 112, E1530–9 (2015).

42. Yuan, M. et al. HMEJ-based safe-harbor genome editing enables efficient generation of cattle with increased resistance to tuberculosis. J. Biol. Chem. 296, 100497 (2021).

43. Gao, Y. et al. Single Cas9 nickase induced generation of NRAMP1 knockin cattle with reduced off-target effects. Genome Biol. 18, 13 (2017).

44. Shanthalingam, S. et al. Precise gene editing paves the way for derivation of Mannheimia haemolytica leukotoxin-resistant cattle. Proc. Natl. Acad. Sci. U. S. A. 113, 13186–13190 (2016).

45. Moennig, V. & Yarnall, M. J. The Long Journey to BVD Eradication. Pathogens vol. 10 1292 Preprint at https://doi.org/10.3390/pathogens10101292 (2021).

46. Chase, C. C. L. The impact of BVDV infection on adaptive immunity. Biologicals 41, 52–60 (2013).

47. Grissett, G. P., White, B. J. & Larson, R. L. Structured literature review of responses of cattle to viral and bacterial pathogens causing bovine respiratory disease complex. J. Vet. Intern. Med. 29, 770–780 (2015).

48. Larson, R. L. Bovine Viral Diarrhea Virus–Associated Disease in Feedlot Cattle. Veterinary Clinics of North America: Food Animal Practice vol. 31 367–380 Preprint at https://doi.org/10.1016/j.cvfa.2015.05.007 (2015).

49. Colett, M. S. et al. Molecular cloning and nucleotide sequence of the pestivirus bovine viral diarrhea virus. Virology 165, 191–199 (1988).

50. Jones, L. R., Zandomeni, R. O. & Weber, E. L. A long distance RT-PCR able to amplify the Pestivirus genome. J. Virol. Methods 134, 197–204 (2006).

51. Xue, W., Mattick, D. & Smith, L. Protection from persistent infection with a bovine viral diarrhea virus (BVDV) type 1b strain by a modified-live vaccine containing BVDV types 1a and 2, infectious bovine rhinotracheitis virus, parainfluenza 3 virus and bovine respiratory syncytial virus. Vaccine 29, 4657–4662 (2011).

52. Neill, J. D. et al. Identification of BVDV2b and 2c subgenotypes in the United States: Genetic and antigenic characterization. Virology 528, 19–29 (2019).

