## Supplementary material for "First gene-edited calf with reduced susceptibility to a major viral pathogen": Table S1, Table S2, Table S3, Table S4, Fig. S1, Fig. S2, Fig. S3, Fig. S4, Fig. S5, Fig. S6

**Supplementary Information**

**Table of Contents**…………………………………………………………...……….……..1

**Table S1.** Virus isolates and BVDV-PI serum used in study. ………………….….………3

**Table S2**. Calf health scoring criteria………………………………………….…...……….4

**Table S3.** Summary of clinical impression scores……………………………….……...........5

**Table S4.** Summary of complete blood count (CBC) values from viral challenge study…………………………...………………………………………………………….......6

**Figure S1.** Genomic sequences of MDBK CD46 gene deletion and CD46 $A_{82}LPTFS_{87}$ substitution……………..…………………………………………………..........7

Legend: Panel **a**, screenshot of IGV software showing approximately 50kb of genomic sequence alignment of parent MDBK $G_{82}QVLAL_{87}$ (top), the *CD46* gene deletion line (middle), and the CD46 $A_{82}LPTFS_{87}$ substitution line (bottom). Panel **b**, screenshot of IGV software showing an 81 bp region centered on the six amino acid substitution site.

**Figure S2.** BVDV infection efficiency in CD46-edited MDBK cells …………......……….8

Legend: MDBK, MDBK-CD46Δ, and MDBK-CD46 $A_{82}LPTFS_{87}$ cells were infected with cytopathic (panel **a**) or non-cytopathic (panel **b**) BVDV isolates at a MOI of 2 and infection efficiency was visualized at 20 hpi by IF. Cells were fixed and stained using an anti-BVDV E2 monoclonal antibody and FITC labeled secondary antibody. Nuclei were stained with DAPI to ensure images were taken in regions with complete cell monolayers (not shown). Cells imaged at 10x magnification. Panel **c**, comparison of viral replication kinetics in MDBK, and MDBK-CD46 $A_{82}LPTFS_{87}$, and MDBK-CD46Δ cells. Cells were infected with BVDV at an MOI of 0.1 and collected by freeze thaw at the indicated times post infection. Viral RNA was detected by RT-qPCR and fold change in viral RNA relative to the input sample (0 hours post-infection) was calculated using the delta Ct method. Results represent the mean ± standard deviation (n = 3).

**Figure S3.** Genomic sequences of Gir CD46 $A_{82}LPTFS_{87}$ substitutions …..…………..9

Legend: One edited CD46 $A_{82}LPTFS_{87}$ fetus was spontaneously aborted at 212 days gestation, but had no obvious physical defects

**Figure S4.** Full term Gir calf with CD46 $A_{82}LPTFS_{87}$ substitutions……………………..10

Legend: An edited CD46 $A_{82}LPTFS_{87}$ Gir calf was delivered by cesarean section at full term (285 days) and was born healthy on July 19, 2021.

**Figure S5.** Genomic sequences of Gir CD46 A$_{82}$LPTFS$_{87}$ substitutions……………..….11

Legend: Screenshot of IGV software showing approximately 154 bp of genomic sequence alignment of the parent Gir cell line G$_{82}$QVLAL$_{87}$ (top panel), the unedited fetus cloned from the same cell line G$_{82}$QVLAL$_{87}$ (2nd panel), the edited cloned fetus with the CD46 A$_{82}$LPTFS$_{87}$ substitution (3rd panel), and the live cloned calf with the CD46 A$_{82}$LPTFS$_{87}$ substitution (4th panel).

**Figure S6.** Genotyping virus from BVDV-PI calf used in challenge study………….……..12

Legend: Panel **a**, to determine whether the BVDV-PI calf was shedding infectious (replication competent) virus, serum (100 uL) or medium from a nasal swab sample (50 ul) was inoculated on MDBK cells for 2 hours at 37°C. Unbound virus was removed by washing cells four times with PBS. Cells were fixed and stained at 72 hpi using an anti-BVDV E2 monoclonal antibody and FITC labeled secondary antibody. Nuclei were stained with DAPI (blue). Panel **b**, RNA was extracted from serum collected from the BVDV-PI calf and sequenced on the Illumina platform. Reads were trimmed and mapped to a reference genome then mapped reads were *de novo* assembled. The WGS was aligned with reference genomes using Muscle 3.8.425 as implemented in Geneious v2022.1.1. A maximum likelihood tree was constructed using FastTree v2.1.11 with an optimized Gamma20 likelihood and the generalized time-reversible (GTR) model.[34] Isolate USMARC_PI20734 (Genbank accession OP548503), marked by a star, is the isolate from the PI calf used in this study.

**Table S1**. Virus isolates and BVDV-PI serum used in study.

| Isolate | Genotype | Biotype | Year | Location | GenBank Accession No. | Reference |
|---|---|---|---|---|---|---|
| NADL | 1a | cp | 1962 | USA | M31182 | 48 |
| Singer | 1a | cp | 1974 | USA | DQ088995 | 49 |
| MDA280n | 1b | cp | NA | USA | NA | NA |
| IIICPE | 1b | cp | 1981 | USA | NA | 50 |
| 296c | 2 | cp | 1995 | USA | MH806436 | 51 |
| 53637 | 2 | cp | 2004 | USA | MH231127 | 51 |
| PI-88-14 USMARC-55477 | 1a | ncp | 2014 | USA:OK | KP941586 | 34 |
| PI-90-2021 | 1a | ncp | 2021 | USA:MO | NA | This study |
| PI-84-2021 | 1b | ncp | 2021 | USA:MO | NA | This study |
| PI-85-2021 | 1b | ncp | 2021 | USA:MO | NA | This study |
| PI-86-2021 | 1b | ncp | 2021 | USA:MO | NA | This study |
| PI-87-2021 | 1b | ncp | 2021 | USA:MO | NA | This study |
| PI-88-2021 | 1b | ncp | 2021 | USA:MO | NA | This study |
| PI-89-2021 | 1b | ncp | 2021 | USA:MO | NA | This study |
| PI-91-2021 | 1b | ncp | 2021 | USA:MO | NA | This study |
| PI-92-2021 | 1b | ncp | 2021 | USA:MO | NA | This study |
| USMARC_PI20734 (PI challenge calf) | 1b | ncp | 2022 | USA:IA | OP548503 | This study |
| PI-92-2014 USMARC-55476 | 2 | ncp | 2014 | USA:OK | KP941585 | 34 |
| PI-65-2014 USMARC-60780 | 2 | ncp | 2014 | USA:OK | KT832823 | 34 |
| PI-15-2014 USMARC-60779 | 2 | ncp | 2014 | USA | KT832822 | 34 |
| PI-83-2014 USMARC-60768 | 2 | ncp | 2014 | USA:TX | KT832821 | 34 |

Abbreviations: cp, cytopathic; ncp, non-cytopathic; NA, not available

**Table S2. Calf health scoring criteria.** A clinical impression score was determined for the calves on days -4 to 21 of the challenge study.

| Points | Attitude | Cough | Nasal Discharge | Eyes | Feces | Rectal Temp (°F) |
|---|---|---|---|---|---|---|
| 0 | bright, alert and responsive with normal appetite | not coughing | normal/clear | normal | normal | 100-100.9 |
| 1 | quiet but alert and responsive | occasionally coughs | small amount of cloudy discharge from one nostril | small amount of ocular discharge from one or both eyes | semi-formed/ pasty | 101-101.9 |
| 2 | depressed, not eating | repeated coughing | cloudy or excessive discharge from both nostrils | moderate amount of ocular discharge from both eyes | loose but stays on top of tender foot | 102-102.9 |
| 3 | - | - | copious discharge from both nostrils | heavy ocular discharge | watery, falls through tender foot without washing | ≥103 |

### Table S3. Summary of clinical impression scores.

**CD46 ALPTFS edited Gir calf**

| Study day | Rectal Temp | Temp Score | Attitude | Attidtude Score | Cough | Cough Score | Nasal Discharge | Nasal Discharge Score | Eyes | Eye Score | Fecal | Fecal Score | Overall Health Score | Comments |
|---|---|---|---|---|---|---|---|---|---|---|---|---|---|---|
| neg 4 | 101.5 | 1 | Bright Alert and Responsive, normal appetite | 0 | Not coughing | 0 | Normal/Clear | 0 | Normal | 0 | Normal | 0 | **1** | |
| neg 3 | 102 | 2 | Bright Alert and Responsive, normal appetite | 0 | Not coughing | 0 | Normal/Clear | 0 | Normal | 0 | Normal | 0 | **2** | |
| neg 2 | 100 | 0 | Bright Alert and Responsive, normal appetite | 0 | Not coughing | 0 | Normal/Clear | 0 | Normal | 0 | Normal | 0 | **0** | |
| neg 1 | 101.6 | 1 | Bright Alert and Responsive, normal appetite | 0 | Not coughing | 0 | Normal/Clear | 0 | Normal | 0 | Normal | 0 | **1** | |
| 0 | 101.3 | 1 | Bright Alert and Responsive, normal appetite | 0 | Not coughing | 0 | Normal/Clear | 0 | Normal | 0 | Normal | 0 | **1** | |
| 1 | 100.3 | 0 | Bright Alert and Responsive, normal appetite | 0 | Not coughing | 0 | Normal/Clear | 0 | Normal | 0 | Normal | 0 | **0** | |
| 2 | 102.8 | 2 | Bright Alert and Responsive, normal appetite | 0 | Not coughing | 0 | Normal/Clear | 0 | Normal | 0 | Normal | 0 | **2** | |
| 3 | 103.5 | 3 | Bright Alert and Responsive, normal appetite | 0 | Not coughing | 0 | Normal/Clear | 0 | Normal | 0 | Normal | 0 | **3** | |
| 4 | 105.6 | 3 | Bright Alert and Responsive, normal appetite | 0 | Not coughing | 0 | Normal/Clear | 0 | Normal | 0 | Normal | 0 | **3** | |
| 5 | 101.5 | 1 | Bright Alert and Responsive, normal appetite | 0 | Not coughing | 0 | Normal/Clear | 0 | Normal | 0 | Normal | 0 | **1** | |
| 6 | 101.8 | 1 | Bright Alert and Responsive, normal appetite | 0 | Not coughing | 0 | Normal/Clear | 0 | Normal | 0 | Normal | 0 | **1** | |
| 7 | 101.6 | 1 | Bright Alert and Responsive, normal appetite | 0 | Not coughing | 0 | Normal/Clear | 0 | Normal | 0 | Normal | 0 | **1** | |
| 8 | 100.5 | 0 | Bright Alert and Responsive, normal appetite | 0 | Not coughing | 0 | Normal/Clear | 0 | Normal | 0 | Normal | 0 | **0** | |
| 9 | 101.3 | 1 | Bright Alert and Responsive, normal appetite | 0 | Not coughing | 0 | Normal/Clear | 0 | Normal | 0 | Normal | 0 | **1** | |
| 10 | 101.6 | 1 | Bright Alert and Responsive, normal appetite | 0 | Not coughing | 0 | Normal/Clear | 0 | Normal | 0 | Normal | 0 | **1** | |
| 11 | 101.5 | 1 | Bright Alert and Responsive, normal appetite | 0 | Not coughing | 0 | Normal/Clear | 0 | Normal | 0 | Normal | 0 | **1** | |
| 12 | 101.1 | 1 | Bright Alert and Responsive, normal appetite | 0 | Not coughing | 0 | Normal/Clear | 0 | Normal | 0 | Normal | 0 | **1** | |
| 13 | 101.2 | 1 | Bright Alert and Responsive, normal appetite | 0 | Not coughing | 0 | Normal/Clear | 0 | Normal | 0 | Normal | 0 | **1** | |
| 14 | 101.4 | 1 | Bright Alert and Responsive, normal appetite | 0 | Not coughing | 0 | Normal/Clear | 0 | Normal | 0 | Normal | 0 | **1** | |
| 15 | 100.6 | 0 | Bright Alert and Responsive, normal appetite | 0 | Not coughing | 0 | Normal/Clear | 0 | Normal | 0 | Normal | 0 | **0** | |
| 16 | 101.4 | 1 | Bright Alert and Responsive, normal appetite | 0 | Not coughing | 0 | Normal/Clear | 0 | Normal | 0 | Normal | 0 | **1** | |
| 17 | 101.7 | 1 | Bright Alert and Responsive, normal appetite | 0 | Not coughing | 0 | Normal/Clear | 0 | Normal | 0 | Normal | 0 | **1** | |
| 18 | 101.8 | 1 | Quiet but alert and responsive, +/- appetite loss | 1 | Not coughing | 0 | Normal/Clear | 0 | Normal | 0 | Normal | 0 | **2** | |
| 19 | 101.2 | 1 | Bright Alert and Responsive, normal appetite | 0 | Not coughing | 0 | Normal/Clear | 0 | Normal | 0 | Normal | 0 | **1** | |
| 20 | 101.3 | 1 | Bright Alert and Responsive, normal appetite | 0 | Not coughing | 0 | Normal/Clear | 0 | Normal | 0 | Normal | 0 | **1** | |
| 21 | 101.3 | 1 | Bright Alert and Responsive, normal appetite | 0 | Not coughing | 0 | Normal/Clear | 0 | Normal | 0 | Normal | 0 | **1** | |

**CD46 WT control Holstein calf**

| Study day | Rectal Temp | Temp Score | Attitude | Attidtude Score | Cough | Cough Score | Nasal Discharge | Nasal Discharge Score | Eyes | Eye Score | Fecal | Fecal Score | Overall Health Score | Comments |
|---|---|---|---|---|---|---|---|---|---|---|---|---|---|---|
| neg 4 | 101.7 | 1 | Bright Alert and Responsive, normal appetite | 0 | Not coughing | 0 | Normal/Clear | 0 | Normal | 0 | Normal | 0 | **1** | |
| neg 3 | 101.3 | 1 | Bright Alert and Responsive, normal appetite | 0 | Not coughing | 0 | Normal/Clear | 0 | Normal | 0 | Normal | 0 | **1** | |
| neg 2 | 100.6 | 0 | Bright Alert and Responsive, normal appetite | 0 | Not coughing | 0 | Normal/Clear | 0 | Normal | 0 | Normal | 0 | **0** | |
| neg 1 | 100.8 | 0 | Bright Alert and Responsive, normal appetite | 0 | Not coughing | 0 | Normal/Clear | 0 | Normal | 0 | Normal | 0 | **0** | |
| 0 | 101.7 | 1 | Bright Alert and Responsive, normal appetite | 0 | Not coughing | 0 | Normal/Clear | 0 | Normal | 0 | Normal | 0 | **1** | |
| 1 | 101 | 1 | Bright Alert and Responsive, normal appetite | 0 | Not coughing | 0 | Normal/Clear | 0 | Normal | 0 | Normal | 0 | **1** | |
| 2 | 102.3 | 2 | Bright Alert and Responsive, normal appetite | 0 | Not coughing | 0 | Normal/Clear | 0 | Normal | 0 | Normal | 0 | **2** | |
| 3 | 104.5 | 3 | Quiet but alert and responsive, +/- appetite loss | 1 | Not coughing | 0 | Small amount of cloudy discharge | 1 | Normal | 0 | Normal | 0 | **5** | Slight, cloudy discharge from left nostril mostly; Feces are not as formed and softer than normal but not quite "pasty" |
| 4 | 102.5 | 2 | Bright Alert and Responsive, normal appetite | 0 | Not coughing | 0 | Normal/Clear | 0 | Normal | 0 | Normal | 0 | **2** | |
| 5 | 103.7 | 3 | Bright Alert and Responsive, normal appetite | 0 | Not coughing | 0 | Normal/Clear | 0 | Normal | 0 | Normal | 0 | **3** | |
| 6 | 100.6 | 0 | Bright Alert and Responsive, normal appetite | 0 | Not coughing | 0 | Normal/Clear | 0 | Normal | 0 | Normal | 0 | **0** | |
| 7 | 101.8 | 1 | Bright Alert and Responsive, normal appetite | 0 | Not coughing | 0 | Normal/Clear | 0 | Normal | 0 | Normal | 0 | **1** | |
| 8 | 101.2 | 1 | Quiet but alert and responsive, +/- appetite loss | 1 | Not coughing | 0 | Normal/Clear | 0 | Normal | 0 | Normal | 0 | **2** | Seemed slightly subdued/lethargic today. Was still interested in food. |
| 9 | 102 | 2 | Bright Alert and Responsive, normal appetite | 0 | Repeated coughing | 2 | Normal/Clear | 0 | Normal | 0 | Normal | 0 | **4** | Rhinitis: Red, inflammed skin around bridge of nose and edges of nostrils, no discharge and coughing |
| 10 | 102 | 2 | Bright Alert and Responsive, normal appetite | 0 | Not coughing | 0 | Normal/Clear | 0 | Normal | 0 | Normal | 0 | **2** | Rhinitis: Red, inflammed skin around bridge of nose and edges of nostrils, no discharge |
| 11 | 100.1 | 0 | Bright Alert and Responsive, normal appetite | 0 | Not coughing | 0 | Small amount of cloudy discharge | 1 | Normal | 0 | Normal | 0 | **1** | Rhinitis: Red, inflammed skin around bridge of nose and edges of nostrils, minimal discharge |
| 12 | 100.8 | 0 | Bright Alert and Responsive, normal appetite | 0 | Not coughing | 0 | Normal/Clear | 0 | Normal | 0 | Normal | 0 | **0** | Rhinitis: Red, inflammed skin around bridge of nose and edges of nostrils, minimal discharge |
| 13 | 100.3 | 0 | Bright Alert and Responsive, normal appetite | 0 | Not coughing | 0 | Normal/Clear | 0 | Normal | 0 | Normal | 0 | **0** | Rhinitis: Red, inflammed skin around bridge of nose and edges of nostrils, minimal discharge |
| 14 | 101.8 | 1 | Bright Alert and Responsive, normal appetite | 0 | Not coughing | 0 | Normal/Clear | 0 | Small amount of discharge | 1 | Normal | 0 | **2** | Nose showing signs of healing |
| 15 | 101.8 | 1 | Bright Alert and Responsive, normal appetite | 0 | Not coughing | 0 | Normal/Clear | 0 | Normal | 0 | Normal | 0 | **1** | |
| 16 | 101.6 | 1 | Bright Alert and Responsive, normal appetite | 0 | Not coughing | 0 | Normal/Clear | 0 | Normal | 0 | Normal | 0 | **1** | |
| 17 | 102 | 3 | Bright Alert and Responsive, normal appetite | 0 | Not coughing | 0 | Normal/Clear | 0 | Normal | 0 | Normal | 0 | **3** | |
| 18 | 101.8 | 1 | Quiet but alert and responsive, +/- appetite loss | 1 | Not coughing | 0 | Normal/Clear | 0 | Normal | 0 | Normal | 0 | **2** | |
| 19 | 100.3 | 0 | Quiet but alert and responsive, +/- appetite loss | 1 | Not coughing | 0 | Normal/Clear | 0 | Normal | 0 | Normal | 0 | **1** | |
| 20 | 101.1 | 1 | Bright Alert and Responsive, normal appetite | 0 | Not coughing | 0 | Normal/Clear | 0 | Normal | 0 | Normal | 0 | **1** | |
| 21 | 101 | 1 | Bright Alert and Responsive, normal appetite | 0 | Not coughing | 0 | Normal/Clear | 0 | Normal | 0 | Normal | 0 | **1** | |

Days in grey are samples taken during the exposure period to the BVDV-PI calf; day of study is relative to the introduction of the BVDV-PI calf. BVDV-PI calf was added to the room after the day 0 samples were collected and was removed at the time of day 7 sample collection.

**Table S4. Summary of complete blood count (CBC) values from viral challenge study.**

**CD46 ALPTFS edited calf**

| Day of Study | WBC | NEU | LYM | MONO | EOS | BAS | NEU % | LYM % | MONO % | EOS % | BAS % | RBC | HGB | HCT | MCV | MCH | MCHC | RDW % | PLT | MPV |
|---|---|---|---|---|---|---|---|---|---|---|---|---|---|---|---|---|---|---|---|---|
| -5 | 6.38 | 1.49 | 4.8 | 0.06 | 0.02 | 0.01 | 23.4 | 75.2 | 0.8 | 0.4 | 0.2 | 7.74 | 10.3 | 31.2 | 40.3 | 13.4 | 33.1 | 21.1 | 370 | 4.3 |
| 0 | 7.06 | 1.57 | 5.29 | 0.12 | 0.07 | 0.01 | 22.2 | 74.9 | 1.8 | 1 | 0.1 | 7.98 | 10.7 | 32.7 | 41 | 13.4 | 32.7 | 21.7 | 457 | 4.2 |
| 2 | 10.24 | 6.25 | 3.83 | 0.13 | 0.03 | 0 | 61 | 37.4 | 1.3 | 0.3 | 0 | 7.09 | 9.5 | 29 | 40.9 | 13.4 | 32.7 | 21.2 | 323 | 4.1 |
| 4 | 4.34 | 1.67 | 2.43 | 0.23 | 0.01 | 0 | 38.5 | 56 | 5.1 | 0.4 | 0 | 6.59 | 8.9 | 26.8 | 40.6 | 13.6 | 33.4 | 21.6 | 326 | 4.1 |
| 7 | 5.81 | 1.24 | 4.33 | 0.19 | 0.04 | 0.01 | 21.3 | 74.5 | 3.3 | 0.6 | 0.3 | 7.6 | 10 | 30.9 | 40.6 | 13.2 | 32.5 | 21.4 | 491 | 4.2 |
| 9 | 7.59 | 1.95 | 5.42 | 0.16 | 0.04 | 0.02 | 25.7 | 71.4 | 2.2 | 0.5 | 0.2 | 7.27 | 9.3 | 29.7 | 40.8 | 12.8 | 31.3 | 21.3 | 574 | 4.3 |
| 11 | 7.15 | 1.76 | 5.19 | 0.14 | 0.05 | 0.01 | 24.6 | 72.6 | 2 | 0.7 | 0.1 | 7.39 | 9.6 | 29.6 | 40.1 | 13.1 | 32.6 | 21.3 | 556 | 4.2 |
| 14 | 8.13 | 2.51 | 5.45 | 0.12 | 0.05 | 0 | 30.9 | 67 | 1.4 | 0.6 | 0.1 | 7.09 | 9.6 | 28.2 | 39.8 | 13.6 | 34.1 | 21.1 | 475 | 4.5 |
| 16 | 8.4 | 2.47 | 5.72 | 0.16 | 0.04 | 0.01 | 29.4 | 68.1 | 1.9 | 0.5 | 0.1 | 7.06 | 9 | 28.1 | 39.8 | 12.8 | 32.1 | 21.6 | 472 | 4.5 |
| 18 | 7.46 | 2.33 | 4.9 | 0.14 | 0.08 | 0.01 | 31.3 | 65.6 | 1.8 | 1.1 | 0.2 | 7.98 | 10.3 | 31.8 | 39.9 | 12.9 | 32.4 | 21.7 | 452 | 4.9 |
| 22 | 7.75 | 2 | 5.54 | 0.11 | 0.09 | 0.01 | 25.8 | 71.5 | 1.4 | 1.1 | 0.2 | 7.79 | 10.3 | 31.2 | 40 | 13.2 | 33.1 | 21.9 | 496 | 5 |
| 25 | 7.96 | 2.23 | 5.45 | 0.15 | 0.1 | 0.03 | 28 | 68.5 | 1.9 | 1.2 | 0.4 | 7.85 | 10.4 | 30.9 | 39.3 | 13.3 | 33.7 | 22.1 | 478 | 4.5 |
| 30 | 9.64 | 3.07 | 6.21 | 0.31 | 0.02 | 0.03 | 31.8 | 64.4 | 3.2 | 0.2 | 0.4 | 7.58 | 9.6 | 30 | 39.6 | 12.7 | 32 | 22.2 | 424 | 4.7 |
| 37 | 9.45 | 2.58 | 6.63 | 0.1 | 0.13 | 0.01 | 27.3 | 70.1 | 1.1 | 1.4 | 0.1 | 9.37 | 12.2 | 38.8 | 41.4 | 13 | 31.3 | 21.5 | 407 | 4.4 |
| 44 | 8.95 | 2.25 | 6.5 | 0.07 | 0.1 | 0.03 | 25.1 | 72.7 | 0.8 | 1.1 | 0.3 | 8.6 | 11.2 | 34.9 | 40.6 | 13 | 32.1 | 21.4 | 326 | 4.6 |

**CD46 WT control calf**

| Day of Study | WBC | NEU | LYM | MONO | EOS | BAS | NEU % | LYM % | MONO % | EOS % | BAS % | RBC | HGB | HCT | MCV | MCH | MCHC | RDW % | PLT | MPV |
|---|---|---|---|---|---|---|---|---|---|---|---|---|---|---|---|---|---|---|---|---|
| -5 | 6.78 | 2.16 | 4.37 | 0.13 | 0.11 | 0.01 | 31.9 | 64.5 | 1.9 | 1.5 | 0.2 | 7.5 | 10.7 | 31.8 | 42.4 | 14.3 | 33.7 | 22.5 | 271 | 5.5 |
| 0 | 7.41 | 1.95 | 5.21 | 0.07 | 0.16 | 0.02 | 26.3 | 70.2 | 1 | 2.2 | 0.3 | 8.29 | 11.9 | 34.5 | 41.7 | 14.3 | 34.4 | 22.2 | 323 | 5.8 |
| 2 | 7.27 | 3.46 | 3.77 | 0.03 | 0.01 | 0 | 47.5 | 51.8 | 0.4 | 0.2 | 0.1 | 6.9 | 10.1 | 29.3 | 42.4 | 14.6 | 34.4 | 22 | 226 | 5.9 |
| 4 | 5.22 | 2.48 | 2.55 | 0.17 | 0.02 | 0 | 47.5 | 48.8 | 3.3 | 0.4 | 0 | 6.62 | 9.6 | 27.5 | 41.5 | 14.5 | 35 | 22.1 | 189 | 5.5 |
| 7 | 4.44 | 1.78 | 2.45 | 0.17 | 0.04 | 0 | 40 | 55.1 | 4 | 0.9 | 0 | 6.9 | 9.6 | 28.5 | 41.3 | 14 | 33.8 | 22.3 | 242 | 5.6 |
| 9 | 5.37 | 1.66 | 3.37 | 0.27 | 0.07 | 0 | 30.9 | 62.6 | 5 | 1.4 | 0.1 | 7.05 | 9.9 | 29.5 | 41.7 | 14 | 33.5 | 22 | 296 | 5.5 |
| 11 | 7.04 | 2.99 | 3.7 | 0.21 | 0.11 | 0.03 | 42.4 | 52.6 | 2.8 | 1.7 | 0.5 | 6.64 | 9.7 | 27.6 | 41.5 | 14.7 | 35.3 | 22.4 | 331 | 5.5 |
| 14 | 8.51 | 3.95 | 3.86 | 0.38 | 0.29 | 0.03 | 46.4 | 45.4 | 4.5 | 3.4 | 0.3 | 6.63 | 9.8 | 27.3 | 41.2 | 14.8 | 35.9 | 22.6 | 365 | 5.6 |
| 16 | 10.29 | 5.75 | 4.16 | 0.3 | 0.05 | 0.03 | 55.8 | 40.4 | 3 | 0.5 | 0.3 | 7.03 | 9.8 | 29 | 41.2 | 13.9 | 33.6 | 23.1 | 369 | 5.8 |
| 18 | 10.53 | 6.2 | 3.83 | 0.36 | 0.1 | 0.04 | 58.9 | 36.4 | 3.3 | 1 | 0.4 | 7.07 | 10.1 | 29.3 | 41.5 | 14.3 | 34.4 | 22.6 | 441 | 5.8 |
| 22 | 8.46 | 3.83 | 3.84 | 0.3 | 0.46 | 0.03 | 45.3 | 45.4 | 3.6 | 5.3 | 0.4 | 6.92 | 10.1 | 28.9 | 41.8 | 14.6 | 34.8 | 22.7 | 483 | 5.6 |
| 25 | 8.45 | 3.36 | 4.38 | 0.33 | 0.34 | 0.04 | 39.8 | 51.8 | 3.9 | 4.1 | 0.4 | 6.72 | 9.7 | 27.9 | 41.5 | 14.4 | 34.6 | 22.9 | 407 | 5.8 |
| 30 | 8.83 | 3.83 | 4.5 | 0.43 | 0.04 | 0.03 | 43.3 | 51 | 4.9 | 0.5 | 0.3 | 7.11 | 9.8 | 29.4 | 41.4 | 13.7 | 33.2 | 22.4 | 312 | 6 |
| 37 | 8.41 | 3.4 | 4.41 | 0.17 | 0.43 | 0 | 40.4 | 52.4 | 2 | 5.2 | 0 | 7.59 | 10.6 | 32.6 | 43 | 13.9 | 32.4 | 21.6 | 308 | 6 |
| 44 | 9.41 | 3.89 | 5.08 | 0.13 | 0.29 | 0.02 | 41.4 | 53.9 | 1.4 | 3.1 | 0.2 | 7.43 | 10.9 | 32 | 43 | 14.6 | 34 | 21 | 326 | 5.9 |

Days in grey are samples taken during the exposure period to the BVDV-PI calf; day of study is relative to the introduction of the BVDV-PI calf. BVDV-PI calf was added to the room after the day 0 samples were collected and was removed at the time of day 7 sample collection. Numbers in red are CBC values that fell outside of the expected (normal) range. Abbreviations: WBC, white blood cells (normal range: 4.6 - 23.8 e6/mL); NEU, neutrophils (1.32 - 9.98 e6/mL; 13.2 - 73.1%); LYM, lymphocytes (1.62 - 20.35 e6/mL; 20 - 82.6%); MONO, monocytes (0 - 2.1 e6/mL; 0 - 13.2%); EOS, eosinophils (0 - 1.08 e6/mL; 0 - 8%); BAS, basophils (0 - 0.29 e6/mL; 0 - 1.5%); RBC, red blood cells (4.6 - 8.2 e6/uL); HGB, hemoglobin (7.8 - 13.8 g/dL); HCT, hematocrit (22 - 42%); MCV, mean corpuscular volume (37 - 56 fL); MCH, mean corpuscular hemoglobin (12.5 - 19.8 pg); MCHC, mean corpuscular hemoglobin concentration (30 - 38 g/dL); RDW, red cell distribution width (16.5 - 26.5%); PLT, platelets (100- 720 e6/mL); MPV, mean platelet volume (4.8 - 7.6 fL).

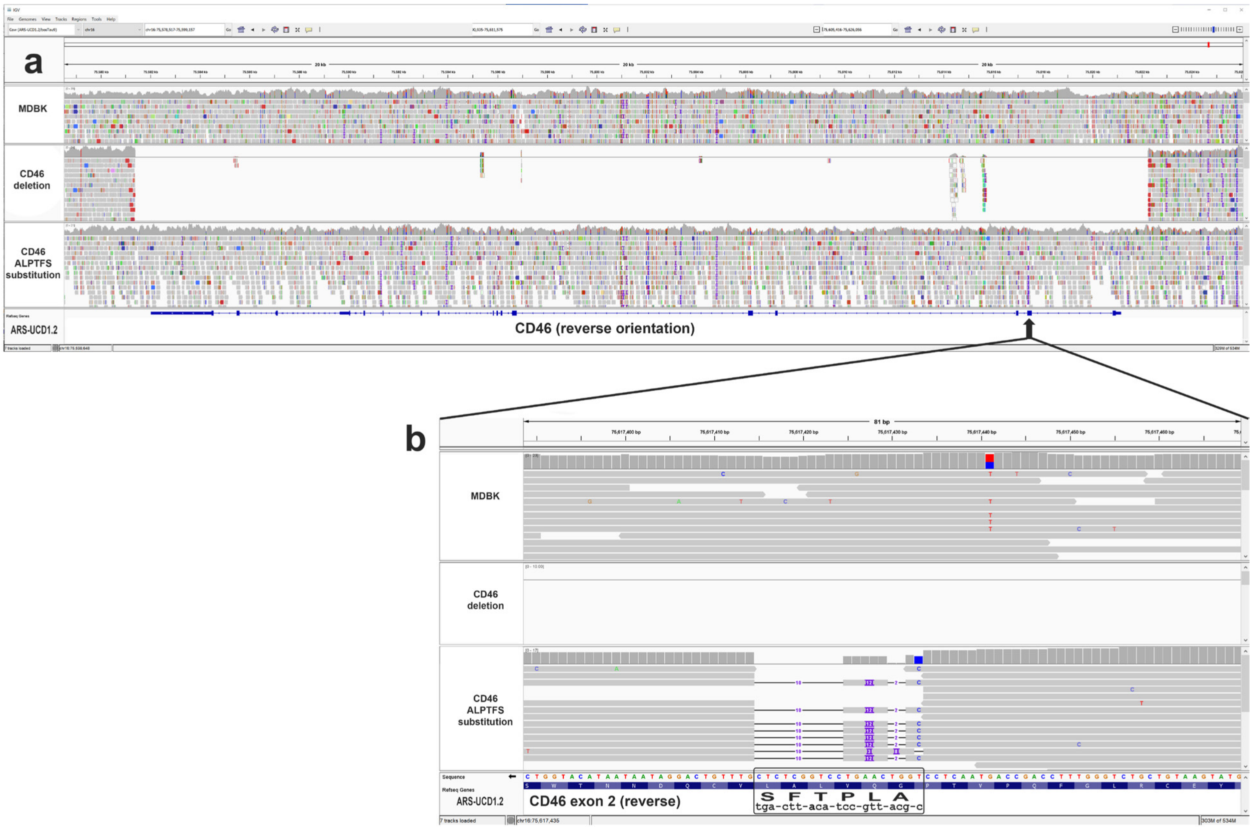


**Figure S1. Genomic sequences of MDBK CD46 gene deletion and CD46 $A_{82}$LPTFS$_{87}$ substitution.** Panel **a**, screenshot of IGV software showing approximately 50kb of genomic sequence alignment of parent MDBK $G_{82}$QVLAL$_{87}$ (top), the *CD46* gene deletion line (middle), and the CD46 $A_{82}$LPTFS$_{87}$ substitution line (bottom). Panel **b**, screenshot of IGV software showing an 81 bp region centered on the six amino acid substitution site.

**MDBK IGV Session URL:** https://s3.us-west-2.amazonaws.com/usmarc.heaton.public/WGS/CellLines/ARS1.2/sessions/ARS12._MDBK_CD46_3tracks.xml

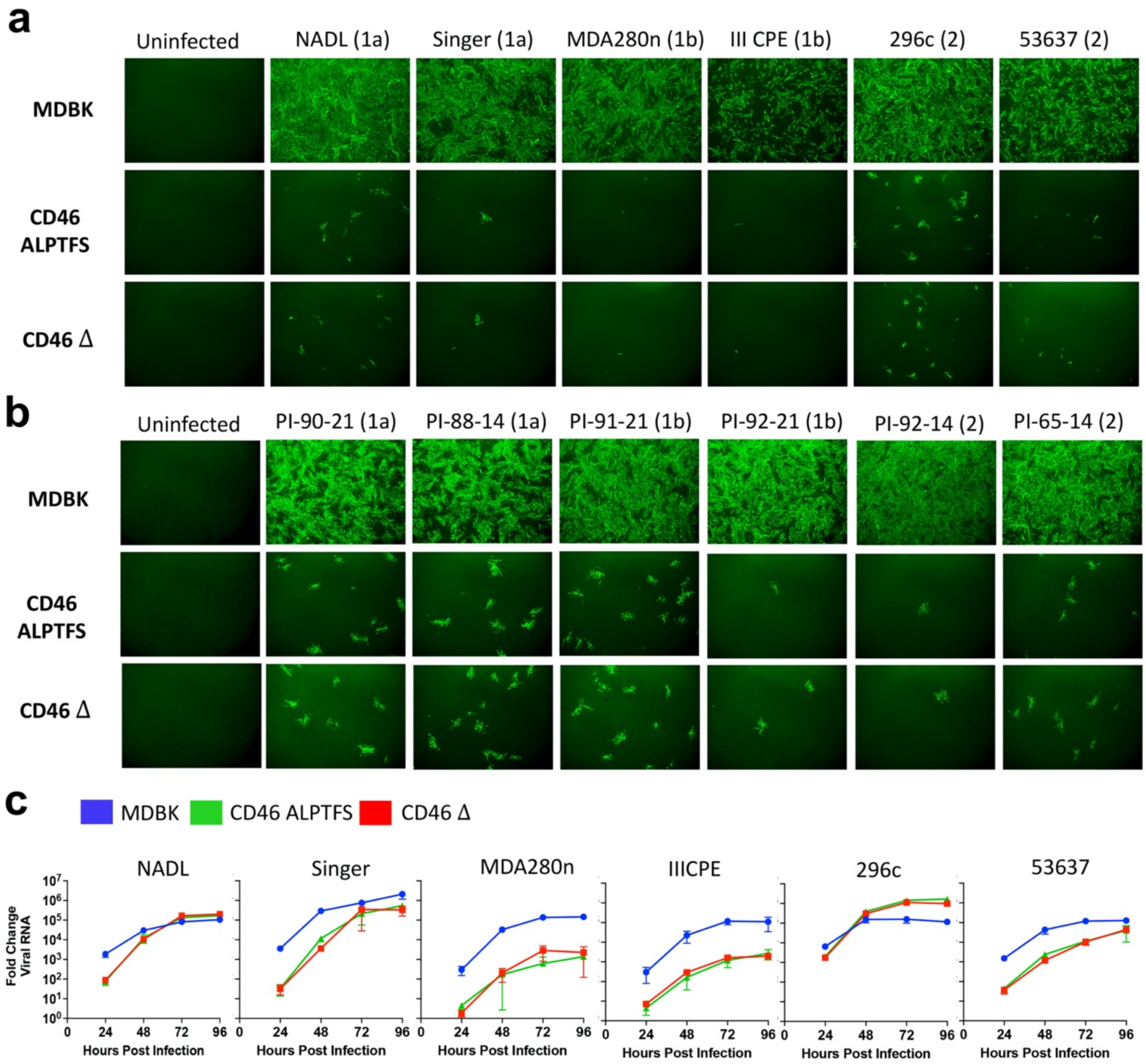


**Figure S2. BVDV infection efficiency in CD46-edited MDBK cells.** MDBK, MDBK-CD46Δ, and MDBK-CD46 $A_{82}LPTFS_{87}$ cells were infected with cytopathic (panel **a**) or non-cytopathic (panel **b**) BVDV isolates at a MOI of 2 and infection efficiency was visualized at 20 hpi by IF. Cells were fixed and stained using an anti-BVDV E2 monoclonal antibody and FITC labeled secondary antibody. Nuclei were stained with DAPI to ensure images were taken in regions with complete cell monolayers (not shown). Cells imaged at 10x magnification. Panel **c**, comparison of viral replication kinetics in MDBK, and MDBK-CD46 $A_{82}LPTFS_{87}$, and MDBK-CD46Δ cells. Cells were infected with BVDV at an MOI of 0.1 and collected by freeze thaw at the indicated times post infection. Viral RNA was detected by RT-qPCR and fold change in viral RNA relative to the input sample (0 hours post-infection) was calculated using the delta Ct method. Results represent the mean ± standard deviation (n = 3).

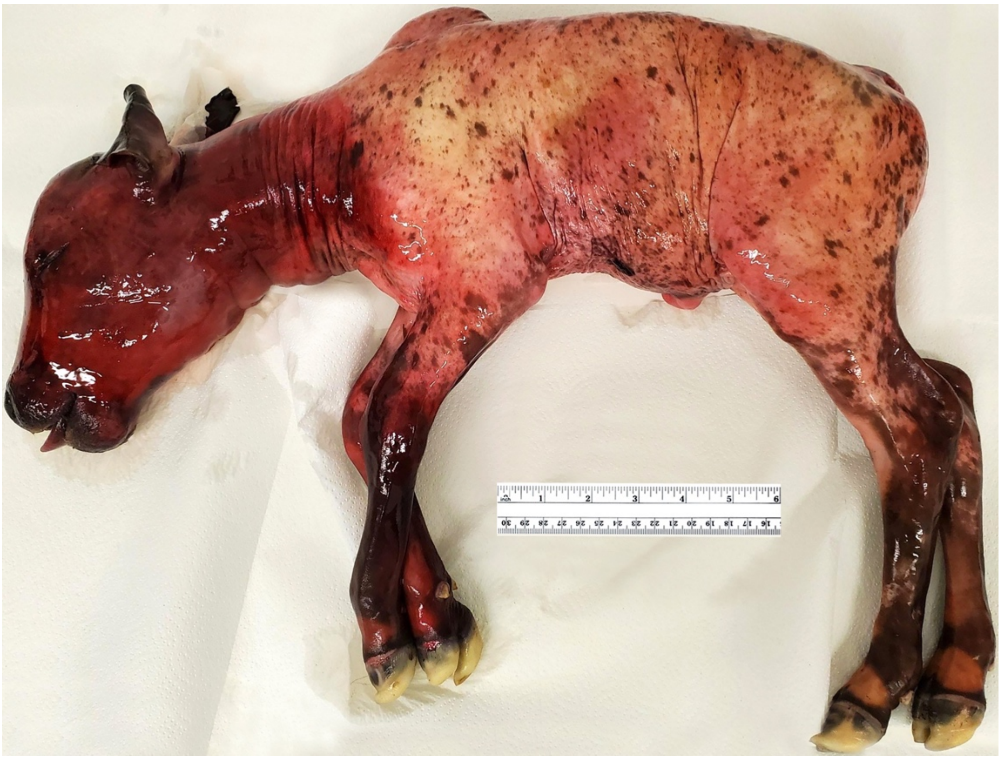

**Figure S3. Aborted Gir calf with CD46 $A_{82}LPTFS_{87}$ substitution.** One edited CD46 $A_{82}LPTFS_{87}$ fetus was spontaneously aborted at 212 days gestation, but had no obvious physical defects.

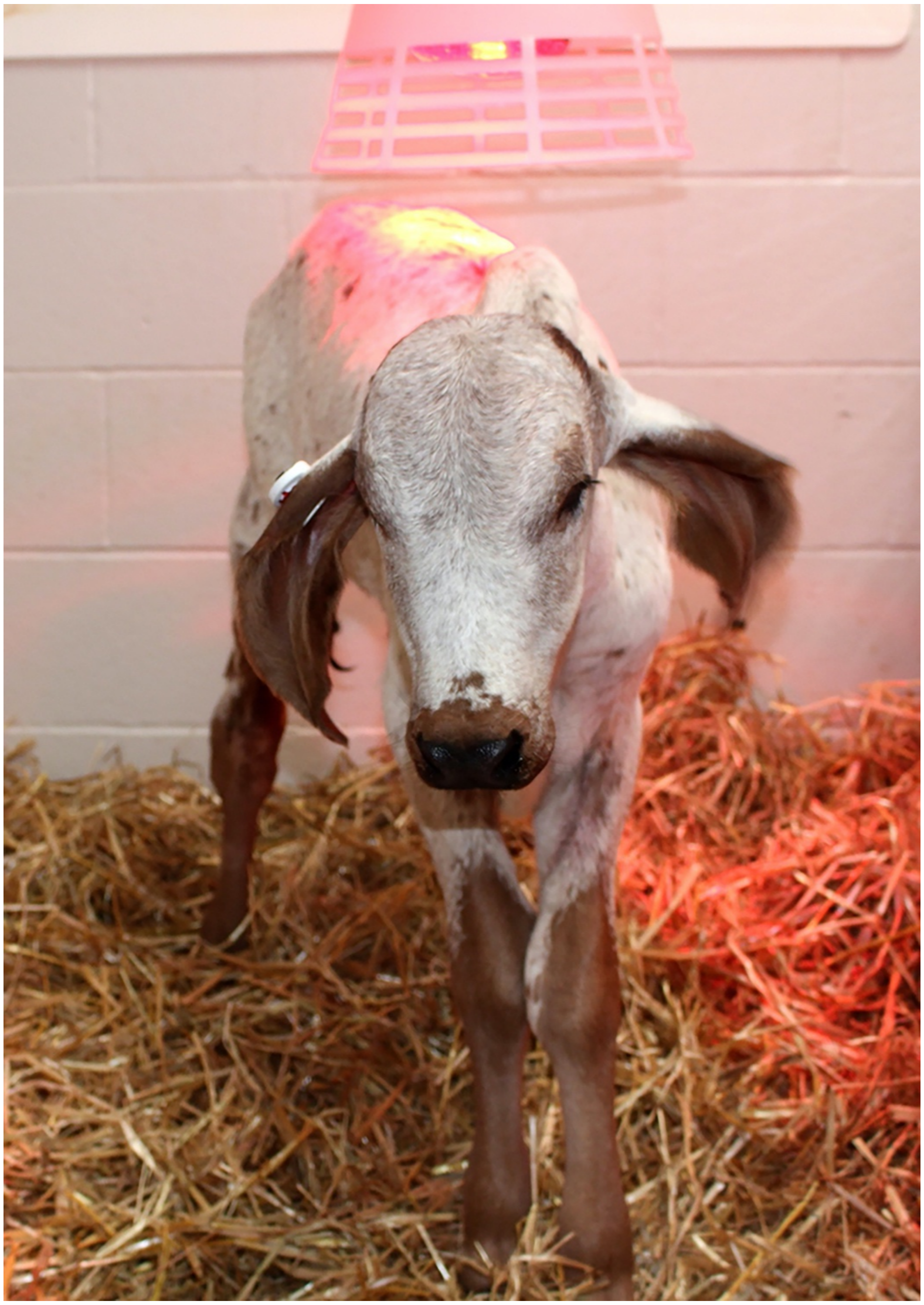

**Figure S4. Full term Gir calf with CD46 $A_{82}LPTFS_{87}$ substitution.** An edited CD46 $A_{82}LPTFS_{87}$ Gir calf (named Ginger) was delivered by cesarean section at full term (285 days) and was born healthy on July 19, 2021.

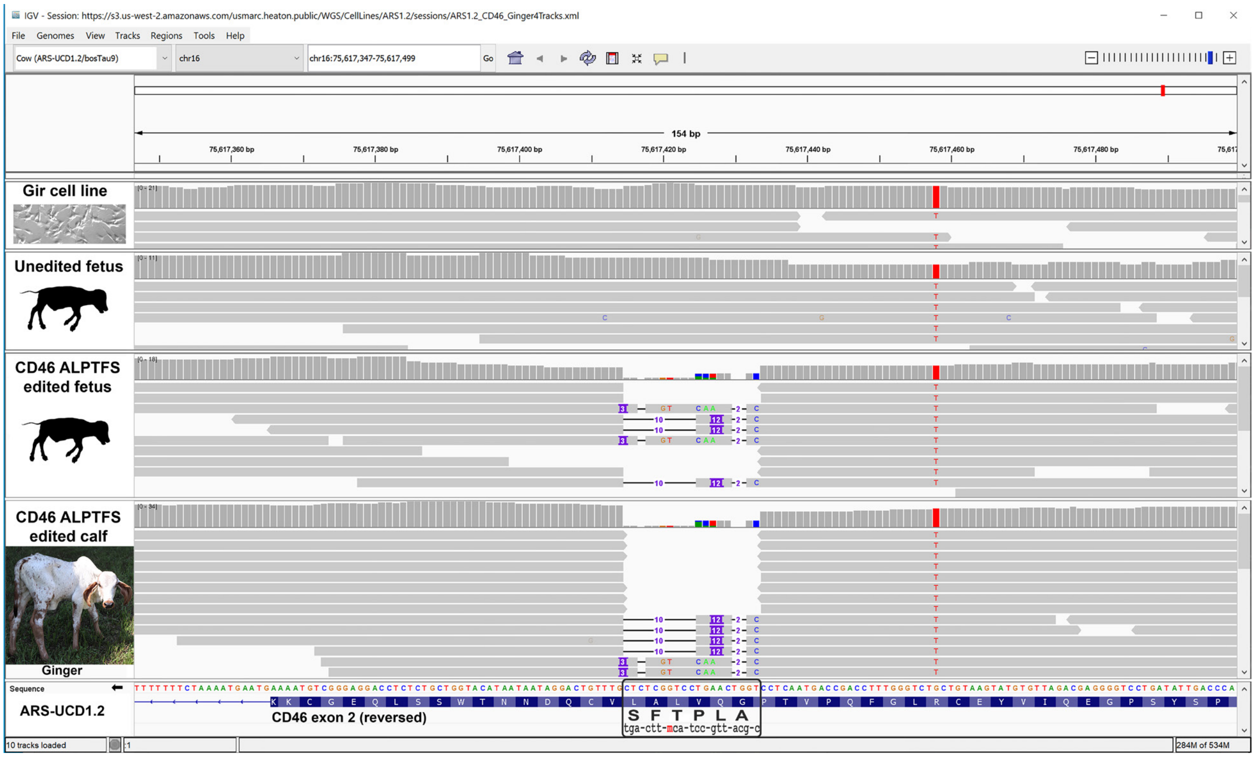


**Figure S5. Genomic sequences of Gir CD46 A$_{82}$LPTFS$_{87}$ substitution.** Screenshot of IGV software showing approximately 154 bp of genomic sequence alignment of the parent Gir cell line G$_{82}$QVLAL$_{87}$ (top panel), the unedited fetus cloned from the same cell line G$_{82}$QVLAL$_{87}$ (2nd panel), the edited cloned fetus with the CD46 A$_{82}$LPTFS$_{87}$ substitution (3rd panel), and the live cloned calf with the CD46 A$_{82}$LPTFS$_{87}$ substitution (4th panel). **Gir clone IGV Session URL**: https://s3.us-west-2.amazonaws.com/usmarc.heaton.public/WGS/CellLines/ARS1.2/sessions/ARS1.2_CD46_Ginger4Tracks.xml

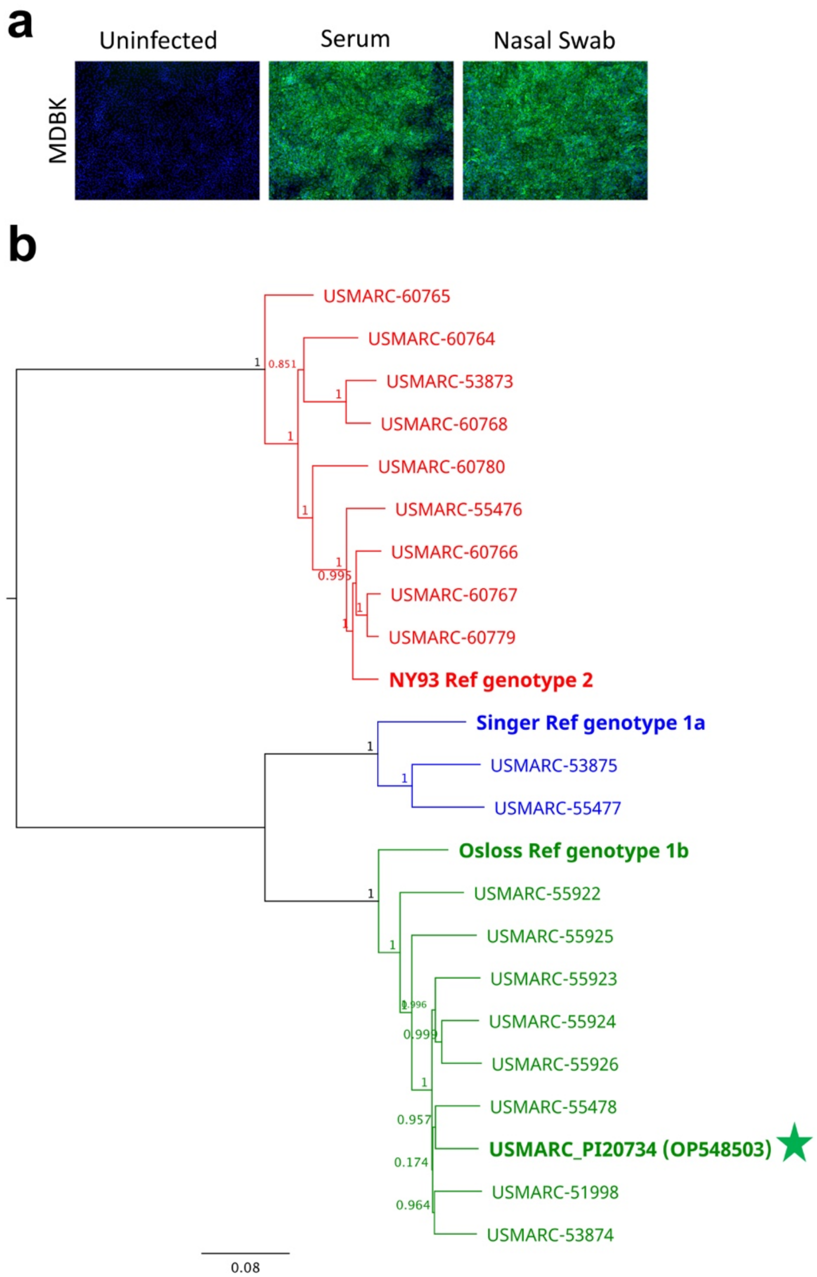


**Figure S6. Genotyping virus from BVDV-PI calf used in challenge study.** Panel **a**, to determine whether the BVDV-PI calf was shedding infectious (replication competent) virus, serum (100 uL) or medium from a nasal swab sample (50 ul) was inoculated on MDBK cells for 2 hours at 37°C. Cells were fixed and stained at 72 hpi using an anti-BVDV E2 monoclonal antibody and FITC labeled secondary antibody. Nuclei were stained with DAPI (blue). Cells imaged at 10x magnification. Panel **b**, RNA was extracted from serum collected from the BVDV-PI calf and sequenced on the Illumina platform. Reads were trimmed and mapped to a reference genome then mapped reads were *de novo* assembled. The WGS was aligned with reference genomes using Muscle 3.8.425 as implemented in Geneious v2022.1.1. A maximum likelihood tree was constructed using FastTree v2.1.11 with an optimized Gamma20 likelihood and the generalized time-reversible (GTR) model[34]. Isolate USMARC_PI20734 (Genbank accession OP548503), marked by a green star, is the isolate from the PI calf used in this study.
